# Diversification of heart progenitor cells by EGF signaling and differential modulation of ETS protein activity

**DOI:** 10.1101/217679

**Authors:** Benjamin Schwarz, Dominik Hollfelder, Katharina Scharf, Leonie Hartmann, Ingolf Reim

**Affiliations:** Friedrich-Alexander University of Erlangen-Nürnberg, Department of Biology, Division of Developmental Biology, Staudtstr. 5, 91058 Erlangen, Germany

## Abstract

For coordinated circulation, vertebrate and invertebrate hearts require stereotyped arrangements of diverse cell populations. This study explores the process of cardiac cell diversification in the *Drosophila* heart, focusing on the two major cardioblast subpopulations: generic working myocardial cells and inflow valve-forming ostial cardioblasts. By screening a large collection of randomly induced mutants we identified several genes involved in cardiac patterning. Further analysis revealed an unexpected, specific requirement of EGF signaling for the specification of generic cardioblasts and a subset of pericardial cells. We demonstrate that the Tbx20 ortholog Midline acts as a direct target of the EGFR effector Pointed to repress ostial fates. Furthermore, we identified Edl/Mae, an antagonist of the ETS factor Pointed, as a novel cardiac regulator crucial for ostial cardioblast specification. Combining these findings we propose a regulatory model in which the balance between activation of Pointed and its inhibition by Edl controls cardioblast subtype-specific gene expression.

## Introduction

The heart consists of a variety of cells with distinct molecular and physiological properties in both vertebrates and invertebrates. A complex regulatory network of transcription factors and signaling pathways orchestrates the specification of these different cell populations and their proper arrangement within a regionalized working myocardium or other functional structures such as valves, inflow and outflow tracts (reviewed in (Greulich, Rudat, & Kispert, 2011; Miquerol & Kelly, 2013; Rana, Christoffels, & Moorman, 2013); for the invertebrate *Drosophila* heart see e.g. (Rolf Bodmer & Frasch, 2010; Lehmacher, Abeln, & Paululat, 2012; Lovato & Cripps, 2016; Reim & Frasch, 2010)). For example, the vertebrate T-box gene *Tbx20* promotes working myocardial fate by restricting *Tbx2* expression and enabling the expression of chamber myocardium-specific genes (Cai et al., 2005; M. K. Singh et al., 2005; Stennard et al., 2005). By contrast, *Tbx2* and *Tbx3* repress working myocardium-specific gene expression and chamber differentiation in the non-chamber myocardium and thus contribute to the formation of endocardial cushions and structures of the conduction system (Christoffels et al., 2004; Hoogaars et al., 2007; R. Singh et al., 2012). Normal endocardial cushion formation also requires COUP-TFII, an orphan nuclear receptor transcription factor that regulates cell fate decisions in several tissues (Lin et al., 2012; S. P. Wu, Yu, Tsai, & Tsai, 2016). In the embryonic mouse myocardium, COUP-TFII is restricted to atrial cardiomyocytes, a pattern consistent with a function in promoting atrial over ventricular fate (Lin et al., 2012; S. P. Wu et al., 2013). This function appears to involve the up-regulation of *Tbx5* (S. P. Wu et al., 2013), another T-box gene with non-uniform cardiac expression and a fundamental role in heart development and human cardiac disease (Basson et al., 1997; Bruneau et al., 1999; Bruneau et al., 2001; Ghosh, Brook, & Wilsdon, 2017; Steimle & Moskowitz, 2017). Furthermore, FGF-mediated receptor tyrosine kinase (RTK) signaling upstream of the cardiogenic transcription factor Nkx2-5 was recently shown to sustain ventricular chamber identity of cardiomyocytes in zebrafish (Pradhan et al., 2017). As emphasized below, spatial restriction of cardiac transcription factors as well as precisely controlled RTK signaling activities are not only important in vertebrate but also invertebrate hearts ((Gajewski, Choi, Kim, & Schulz, 2000; Lo & Frasch, 2001; Zaffran et al., 2006) and this work).

The *Drosophila* heart (dorsal vessel) comprises several types of cardiomyocytes (in the embryo called cardioblasts, CBs) and non-contractile pericardial cells (PCs) (Rolf Bodmer & Frasch, 2010; Lovato & Cripps, 2016). The progenitors of these cells are specified in segmentally repeated heart fields located at the intersection of BMP/Dpp and Wg/Wnt signaling activities (Frasch, 1995; Reim & Frasch, 2005; X. Wu, Golden, & Bodmer, 1995). Subsequent specification of the definitive cardiogenic mesoderm depends on a conserved group of transcription factors, most importantly those encoded by the *Nkx2-5* ortholog *tinman* (*tin*), the *Gata4* ortholog *pannier* (*pnr*) and the *Dorsocross1-3* T-box genes (three *Tbx6*-related paralogs that also share features with *Tbx2/3/5*; in the following collectively called *Doc*) ((Alvarez, Shi, Wilson, & Skeath, 2003; Azpiazu & Frasch, 1993; R. Bodmer, 1993; Gajewski, Fossett, Molkentin, & Schulz, 1999; Junion et al., 2012; Reim & Frasch, 2005; Reim, Lee, & Frasch, 2003); reviewed in (Reim & Frasch, 2010; Reim, Frasch, & Schaub, 2017)).

While the identification of cardiogenic factors has greatly improved our understanding of early specification events, much less is known about the mechanisms that lead to the diversification of cardiac cell subpopulations. In this study, we mainly focus on the development of the two major cardioblast subpopulations: generic cardioblasts (gCBs), which make up the main portion of the contractile tube ("working myocardium"), and ostial cardioblasts (oCBs), which form bi-cellular valves (ostia) for hemolymph inflow. Due to Hox gene inputs, ostial progenitor specification is limited to the abdominal region ((Lo, Skeath, Gajewski, Schulz, & Frasch, 2002; Lovato, Nguyen, Molina, & Cripps, 2002; Ponzielli et al., 2002; Ryan, Hoshizaki, & Cripps, 2005), reviewed in (Monier, Tevy, Perrin, Capovilla, & Semeriva, 2007)). Current research suggests that each abdominal hemisegment generates at least seven distinct progenitors that give rise to six CBs (4 gCBs + 2 oCBs) and several types of PCs (Tin^+^/<u>E</u>ven-skipped(Eve)^+^ EPCs, <u>T</u>in^+^ TPCs, and <u>O</u>dd-skipped(Odd)^+^ OPCs; (Rolf Bodmer & Frasch, 2010) and references therein). Whereas gCBs (a.k.a. Tin-CBs) maintain expression of *tin*, oCBs (a.k.a. Svp-CBs) specifically express the *COUP-TFII* ortholog *seven-up* (*svp*) and *Doc* (Gajewski et al., 2000; Lo & Frasch, 2001; Ward & Skeath, 2000; Zaffran et al., 2006). Previous work has shown that *Doc* is repressed in gCBs in a *tin*-dependent manner (Zaffran et al., 2006). Robust *tin* expression in turn depends on the *Tbx20* ortholog *midline* (*mid/nmr2*). The *mid* gene is first activated in gCB progenitors, but later, like its paralog *H15/nmr1*, becomes expressed in all cardioblasts (Miskolczi-McCallum, Scavetta, Svendsen, Soanes, & Brook, 2005; Qian, Liu, & Bodmer, 2005; Reim, Mohler, & Frasch, 2005). In oCBs, *svp* represses *tin* expression thereby permitting continued *Doc* expression in these cells (Gajewski et al., 2000; Lo & Frasch, 2001; Zaffran et al., 2006). In the abdomen, gCBs and most PCs are preceded by a precursor that undergoes symmetric division, whereas oCBs and half of the OPCs are derived from common, asymmetrically dividing CB/PC progenitors (Alvarez et al., 2003; Han & Bodmer, 2003; Ward & Skeath, 2000).

The process of progenitor specification in the somatic and cardiogenic mesoderm involves the antagonistic actions of RTK/Ras/MAPK and Delta/Notch signaling (Carmena et al., 2002; Grigorian, Mandal, Hakimi, Ortiz, & Hartenstein, 2011; Hartenstein, Rugendorff, Tepass, & Hartenstein, 1992). Two types of RTKs, the fibroblast growth factor (FGF) receptor Heartless (Htl) and the epidermal growth factor (EGF) receptor EGFR, act positively on progenitor selection via MAPK signaling, although they are used by different progenitors to different extents (Buff, Carmena, Gisselbrecht, Jimenez, & Michelson, 1998; Carmena et al., 2002; Michelson, Gisselbrecht, Zhou, Baek, & Buff, 1998). Htl and its FGF8-like ligands Pyramus (Pyr) and Thisbe (Ths) have a dual function as regulators of mesodermal cell migration and cell specification, with progenitors of the Eve^+^ lineage as the most prominent example for the latter (reviewed in (Bae, Trisnadi, Kadam, & Stathopoulos, 2012; Muha & Müller, 2013)). The exact contribution of EGFR signaling to *Drosophila* heart development has been less clear until now, but it was shown that EGFR loss of function results in a severe reduction of the numbers of cardioblasts, pericardial nephrocytes, and blood progenitors (Grigorian et al., 2011). Molecularly, the predominant EGFR ligand in the embryo, Spitz (Spi), relies on the protease Rhomboid (encoded by *rho*) and the chaperon Star (S) for its conversion from a membrane-bound into its active form (reviewed in (Shilo, 2014)). Among the most important downstream effectors of RTK/Ras/MAPK pathways are the ETS transcription factors PntP2 (encoded by *pointed/pnt*) and Yan/Aop (encoded by *anterior open/aop*). While PntP2 becomes an active transcriptional activator upon phosphorylation by MAPK, the transcriptional repressor Yan is negatively regulated by MAPK (Gabay et al., 1996; O'Neill, Rebay, Tjian, & Rubin, 1994). Unlike PntP2, a shorter isoform encoded by *pnt*, PntP1, is constitutively active but was shown to require activated MAPK for its transcriptional activation at least in some cell types (Brunner et al., 1994; Gabay et al., 1996; Klämbt, 1993; O'Neill et al., 1994). Notably, chordate Pnt orthologs (ETS1/2) were shown to contribute to cardiac progenitor formation in the tunicate *Ciona* and during transdifferentiation of human dermal fibroblasts into cardiac progenitors (Davidson, Shi, Beh, Christiaen, & Levine, 2006; Islas et al., 2012). During early *Drosophila* cardiogenesis, Pnt favors expression of *eve* over that of another homeobox gene, *ladybird* (*lbe*, expressed in mesodermal cells immediately anterior of the Eve^+^ cluster and later in TPCs and two of the four gCBs per hemisegment; (K. Jagla et al., 1997)) (Liu et al., 2006). In addition, Pnt promotes pericardial cell development and antagonizes CB fate, especially that of oCBs (Alvarez et al., 2003).

Despite the progress in the understanding of cardiac progenitor specification, the mechanisms that diversify progenitors of the oCB and gCB lineages have remained elusive. We have performed an unbiased large-scale mutagenesis screen to identify genes that regulate cardiac development in *Drosophila* embryos and found several mutants that show CB subtype-specific defects. On this basis we discovered a novel and rather unexpected function of the EGF pathway in specifying the gCBs of the working myocardium, thus revealing an intimate link between cardioblast specification and diversification. Furthermore we identified ETS domain lacking (Edl a.k.a. Modulator of the activity of ETS, Mae) as a crucial regulator of the specification of inflow valve-forming oCBs. Edl possesses a SAM domain, which mediates binding to the SAM domain-containing ETS factors PntP2 and Yan, thereby inhibiting their activity as a transcriptional activator or repressor, respectively (Baker, Mille-Baker, Wainwright, Ish-Horowicz, & Dibb, 2001; Qiao et al., 2006; Qiao et al., 2004; Tootle, Lee, & Rebay, 2003; Vivekanand, Tootle, & Rebay, 2004; Yamada, Okabe, & Hiromi, 2003). Our data imply that Edl enables *svp* expression and thus oCB fate by limiting the activity of PntP2, thereby blocking subsequent activation of important downstream targets such as *pntP1* and *mid*. Collectively, our data provide the basis for an elaborated model of cardiac cell fate diversification that links MAPK signaling, Pnt activity and the cell type-specific expression patterns of key cardiac transcription factors.

## Results

### Novel EMS-induced mutants reveal a specific requirement of EGF signaling for the specification of generic cardioblasts

In order to identify genes involved in heart and muscle development in an unbiased manner we have performed an EMS mutagenesis screen for chromosome 2 in *Drosophila melanogaster* embryos (Hollfelder, Frasch, & Reim, 2014). Several of the isolated mutants display a partial loss or irregular alignment of cardioblasts (CBs). Such defects may potentially result from mutations in genes that regulate the specification or differentiation of all CBs or only a particular CB subtype. In the latter case, disturbances in the characteristic "2+4" CB pattern of two ostial cardioblast (oCBs; Doc^+^/Tin^-^) and four generic CBs (gCBs; Doc^-^/Tin^+^) per hemisegment are to be expected. To analyze the cardiac pattern of mutants in more detail, we performed immunofluorescent double stainings for Doc and H15 (or alternatively Mef2) to label oCBs and all CBs, respectively. We then genetically and in part also molecularly mapped the mutations responsible for CB pattern anomalies (for details see the Materials and Methods section and Table S1). The class of mutants characterized by a loss of CBs contained several novel alleles of genes involved in RTK/Ras/MAPK signaling, which is consistent with the assumed role of this pathway in cardiac progenitor selection or maintenance (Carmena et al., 2002; Grigorian et al., 2011). However, no specific role for the specification of a particular cardioblast subtype or diversification of gCB versus oCB progenitors had been previously attributed to RTK/Ras/MAPK signaling. Our phenotypic analysis now shows that diminished EGF/EGFR but not FGF/Htl signaling leads to a preferential reduction of gCB numbers. Embryos with partially reduced FGF/Htl signaling, i.e. mutants lacking both copies of the FGF-encoding gene *pyr* and one copy of its paralog *ths*, as well as hypomorphic *htl* mutants, show an about equal reduction of gCB and oCB numbers (Figure 1B, for quantification see Figure 1M; additional examples in Figure 1-figure supplement 1B,C). This CB reduction can be sufficiently explained by uneven spreading of the early mesoderm to Dpp-receiving areas. By contrast, several mutations mapped to EGF signaling components feature a preferential loss of gCBs. In strong *Egfr* mutants very few CBs can be found (Figure 1C, Figure 1-figure supplement 1E). Remarkably, the overwhelming majority of the residual CBs express Doc. The few remaining Doc-negative CBs are usually located towards the anterior and thus are possibly remnants of the oCB-free anterior aorta. In *spitz, rhomboid* and *Star* loss-of-function mutants the number of Doc^-^/Tin^+^ CBs is strongly reduced while that of ostial Doc^+^/Tin^-^ CBs is nearly normal or in some cases even increased by a few cells (Figure 1D-G,M, Figure 1-figure supplement 1E,F, Figure 1-figure supplement 2A-C). CBs apparently do not require activity of the ostial marker gene *svp* to develop and survive independently of EGF, since total CB numbers are similar in *Star* single and *Star svp* double mutants (compare Figure 1H to 1G; quantification in Figure 1M).

**Figure 1.**
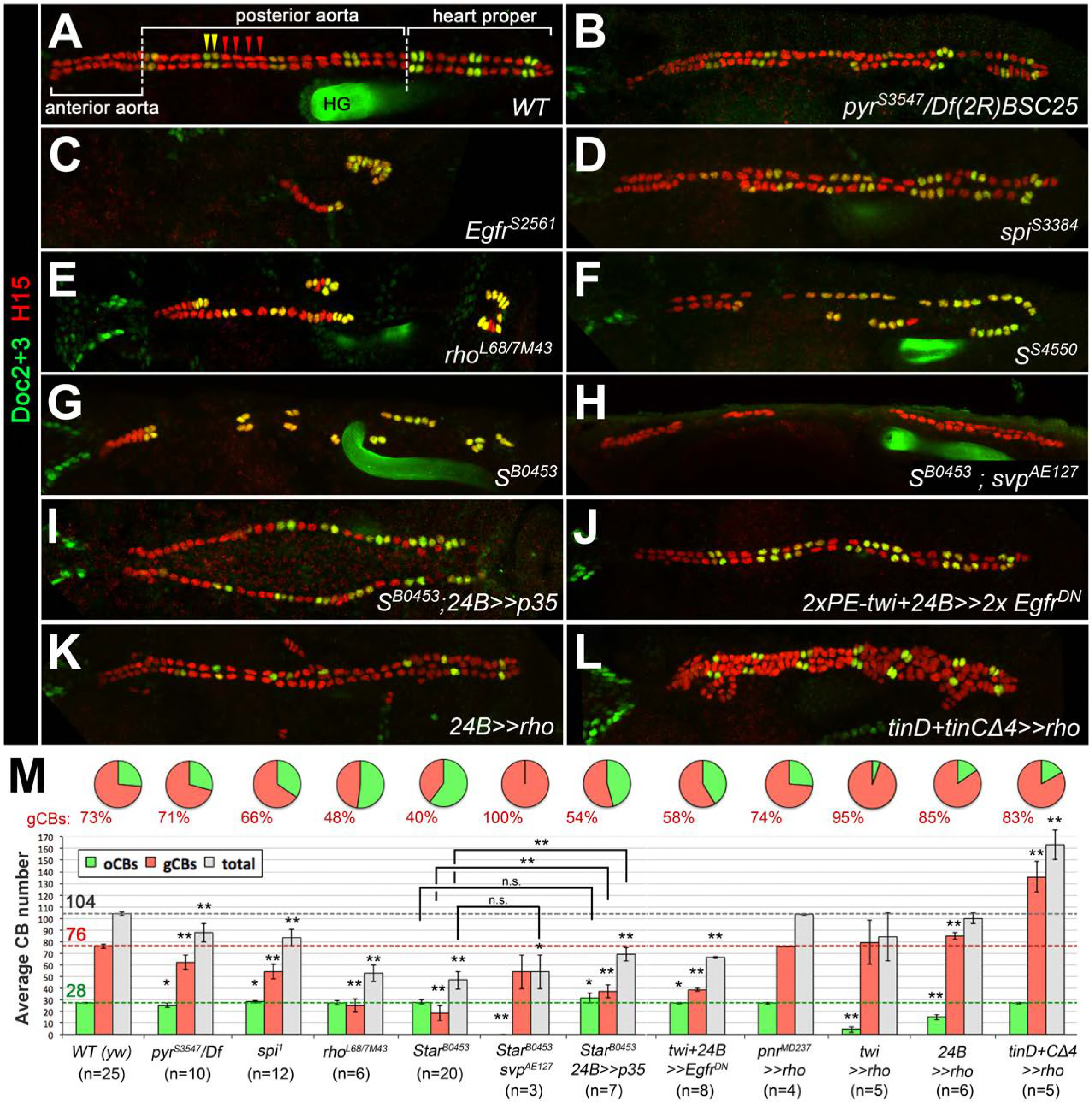
Genetic manipulation of EGF but not FGF signaling leads to cardioblast subtype-specific heart defects. Immunostaining for the cardioblast marker H15 (red) and the ostial cardioblast marker Dorsocross (anti-Doc2+3, green). (HG: hindgut with artificial staining in the lumen). All figures depict dorsal views of stage 16 embryos with anterior to the left unless noted otherwise. (A) Wild type (*WT*) CB pattern with regular alternation of gCBs (red) and oCBs (yellow) in the posterior aorta and the heart proper. The anterior aorta consists entirely of Doc^*-*^ CBs. (B) Mutant with reduced FGF activity (*pyr*^*S3547*^ over a deficiency, *Df(2R)BSC25*, that removes *pyr* and *ths*) showing a reduction of both CB types. (C) Homozygous *Egfr*^*S2561*^ mutant with a severe loss of CBs. Almost all remaining CBs are Doc^+^. Predominant reduction of gCBs is also observed in the EGF pathway-impairing *spitz* group mutants *spi*^*S3384*^ (D), *rho*^*7M43*^/*rho*^*L68*^ (E), *S*^*S4550*^ (F) and *S*^*B0453*^ (G, showing an extreme case in which all retained CBs except for those of the anterior aorta are Doc^+^). (H) In *S* ^*B0453*^ *svp*^*AE127*^ double mutants, total CB numbers are similar to that of *S* single mutants, even though all CBs are Doc-negative. (I) If the apoptosis inhibitor p35 is artificially expressed in the mesoderm of *S* mutants a mild increase in the number of CBs can be observed. Compared to the wild type, more Doc^+^ CBs are present. (J) Pan-mesodermal overexpression of dominant-negative *Egfr* results in a phenotype similar to *spitz* group mutants. Expression of *rho* in the entire mesoderm via *how^24B^-GAL4* (K) or at later time in dorsal mesoderm cells via *tinD+tinCΔ4-GAL4* (L) generates supernumerary gCBs. By contrast, oCB specification is either reduced (K) or unaffected (L) in these backgrounds. (M) Quantification of Doc^+^ oCBs (green), Doc^-^ gCBs (red) and total cardioblasts (grey). The column bar chart depicts average numbers with standard deviation error bars. Asterisks indicate significant differences compared to the *y w* control (*WT*) assessed by Student's t-test (two-tailed, type 3; * = p<0.05, ** = p<0.001; n.s. = not significant). Comparisons between other genotypes are indicated by brackets. Pie charts display the corresponding average fraction of oCBs and gCBs.

Previous studies in EGF pathway mutants suggested that incorrectly specified mesodermal progenitors undergo apoptosis (Buff et al., 1998; Grigorian et al., 2011). Using TUNEL and anti-activated caspase stainings we could not reliably detect signs of apoptosis in the Doc-labeled cardiogenic mesoderm of *Star* mutants, while numerous signals were observed in other tissues (Figure 1-figure supplement 3 and data not shown). If the baculoviral apoptosis inhibitor p35 (Zhou et al., 1997) is artificially expressed in the mesoderm of *S* mutants the number of CBs slightly increases in comparison to *S* mutants without p35 (Figure 1I,M). Although this is consistent with a pro-survival function of EGF signaling, it does not fully account for the gCBs missing in *S* mutants and suggests that the presumptive gCB progenitors largely adopt other fates at reduced EGFR activity. Collectively, these phenotypes imply that EGF signaling plays a major role in the correct specification of gCBs.

### Generic CBs and a subset of Odd^+^ pericardial cells require spatially and temporally coordinated EGF signals

Because EGF signaling is involved in multiple processes during embryogenesis we next asked whether its impact on gCB specification is directly linked to signaling activity within mesoderm cells. Indeed, mesoderm-specific attenuation of the pathway by expression of a dominant-negative EGFR variant resulted in essentially the same phenotype as with the *spitz* group mutants (Figure 1J,M). Activation of the EGF pathway in mesoderm cells appears to be largely controlled by the spatially restricted expression of *rho* (Bidet, Jagla, Da Ponte, Dastugue, & Jagla, 2003; Grigorian et al., 2011; Halfon et al., 2000). In the wild type, *rho* expression is first seen in the Eve^+^ progenitor P2 (Buff et al., 1998; Carmena, Bate, & Jimenez, 1995; Halfon et al., 2000) followed by expression in the adjacent CB progenitor-containing clusters C14 and C16 ((Bidet et al., 2003; Grigorian et al., 2011); see also Figure 1-figure supplement 4A-D). Overexpression of *rho* with the pan-mesodermal *how^24B^-GAL4* driver has been previously reported to affect the number of *tin*-expressing pericardial cells (Bidet et al., 2003), but CBs and their subtypes were not unambiguously labeled in these experiments. We extended these experiments using also other drivers.

Consistent with a mesoderm-autonomous function, overexpression of *rho* in the dorsal ectoderm (via *pnr^MD237^-GAL4*) has no significant effect on CB number or pattern (Figure 1M and data not shown). By contrast, all mesodermal *rho* overexpression setups increase the gCBs:oCBs ratio in comparison to the wild type (Figure 1K-M and data not shown). The impact on the absolute CB numbers depends on the timing and strength of transgene expression. The later *rho* is activated in mesodermal cells (with following drivers according to their temporal order and progressive spatial restriction: *twist-GAL4, how^24B^-GAL4* and *tinD+tinCΔ4-GAL4*) the larger the total number of CBs (Figure 1K-M and data not shown).

Since half of the *odd*-expressing pericardial cells (OPCs) are siblings of oCBs we also analyzed PCs in EGF-related mutants by Odd/Eve double-stainings (Figure 2A-C and data not shown). Consistent with the results of previous studies on Eve^+^ progenitor derivatives (Buff et al., 1998; Carmena et al., 2002; Su, Fujioka, Goto, & Bodmer, 1999), we detected EPCs in almost normal numbers in *spi* group mutants and in embryos with pan-mesodermal dominant-negative EGFR, whereas *spi-*dependent Eve^+^ DA1 muscles were largely absent. OPCs are strongly reduced in these loss-of-function backgrounds. Our quantification revealed that about half of the OPCs were lost in *rho*^*7M43/L68*^ mutants (-45%, n=4). A converse phenotype with many extra OPCs is generated by *rho* overexpression with *tinD+tinCΔ4-GAL4* (Figure 2D). Notably, the number of oCB-sibling OPCs (as identified by *svp-lacZ* reporter analysis) is not significantly reduced in *Star* mutants if compared to the wild type (Figure 2E,F), thus implying that the EGF signaling-dependent OPCs are those derived from symmetrically dividing OPC progenitors.

**Figure 2.**
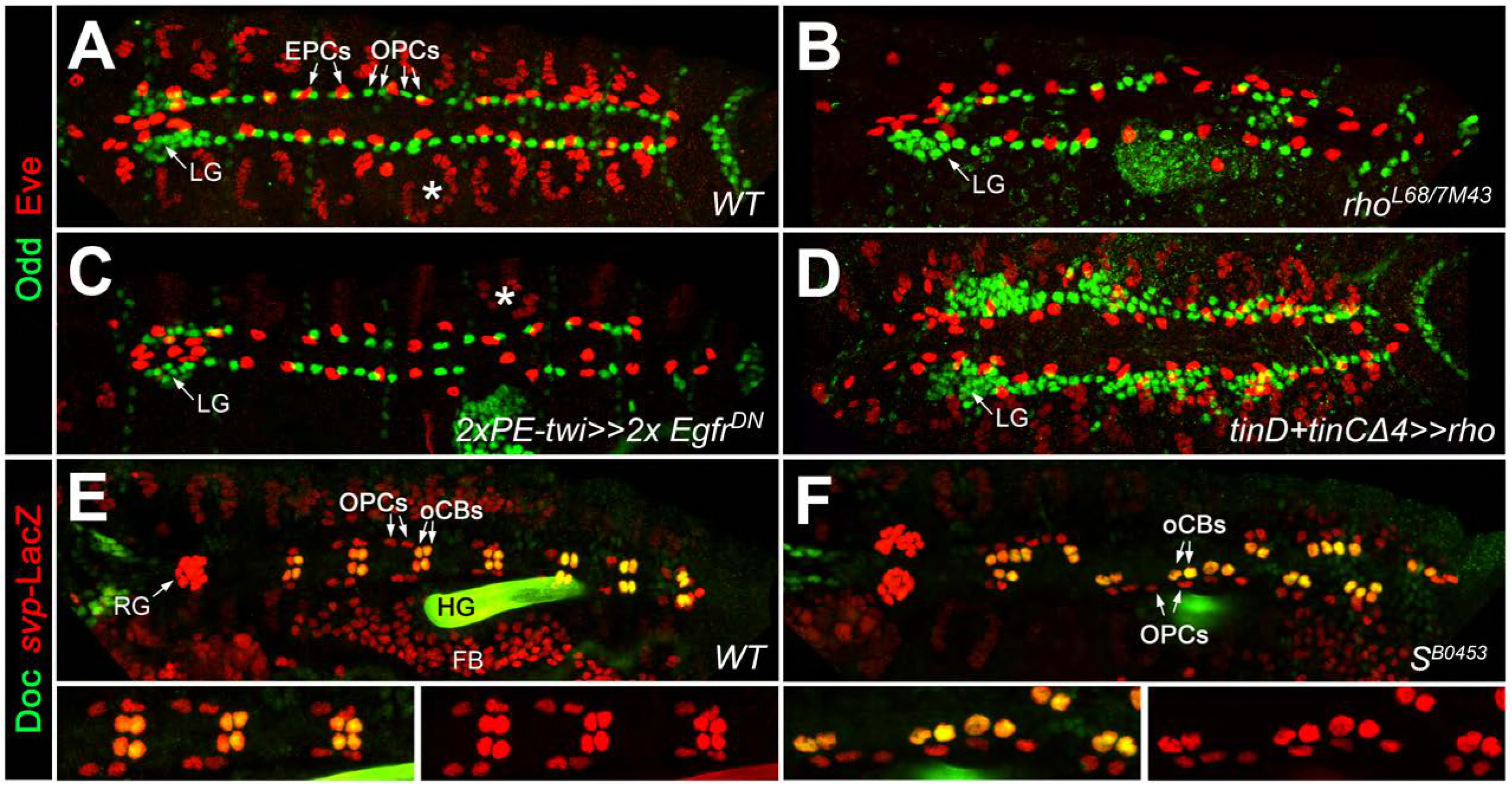
EGF signaling promotes the formation of Odd^+^ PCs. (A-D) Odd/Eve staining to analyze pericardial cells (PCs). (A) In the wild type, each hemisegment contains four OPCs, two EPCs and one Eve^+^ somatic muscle DA1 (*). (B) Amorphic *rho*^*7M43/L68*^ mutant with a loss of about half of all OPCs and all DA1 muscles. (C) Pan-mesodermal overexpression of the dominant-negative *Egfr* results in a phenotype similar to *rho* mutants. (D) Overexpression of *rho* in the dorsal mesoderm generates supernumerary OPCs. (E,F) Doc2+3/β-galactosidase staining in wild type (E) and *Star* mutant embryos (F) carrying a heterozygous copy of *svp^AE127^-lacZ* and showing presence of normal numbers of oCBs (Doc^+^/LacZ^+^) and their OPC siblings (Doc^-^/LacZ^+^). Bottom panels show a higher magnification and β-galactosidase single channel view of the upper panel. LG: lymph gland, RG: ring gland, FB: fat body.

Altogether, these data demonstrate that EGF pathway activity is required in the mesoderm specifically for the specification of the symmetrically dividing gCB and OPCs progenitors, but is largely dispensable or even detrimental for the specification of the *svp*-expressing oCB/OPC progenitors.

### The SAM domain protein Edl promotes specification of ostial cardioblasts by blocking Pointed activity

Our EMS screen also yielded mutants in which the number of ostial cardioblasts was specifically reduced. One such complementation group consisting of three alleles was mapped to the *numb* gene (alleles listed Table S1), which is consistent with its well-known function as a Notch suppressor during asymmetric cell division in the oCB lineage (Gajewski et al., 2000; Ward & Skeath, 2000). Preferential reduction of oCBs was also observed in the mutant line *S0520*. We found that its cardiac phenotype was caused by loss of the gene *ETS domain lacking* (*edl*) as part of a multi-gene deletion and named this mutant *Df(2R)edl-S0520* (Figure 3A, Table S2). We identified *edl* as the gene responsible for the oCB losses by obtaining phenocopies with other *edl* mutants (Figure 3A-D and data not shown). The *lacZ* enhancer trap insertion allele *edl*^*k06602*^ was used in most *edl* loss-of-function experiments since its cardiac phenotype is indistinguishable from that of *Df(2R)edl-S0520* and *Df(2R)edl-L19* (Figure 3C,D and data not shown), and we detected in this strain a small deletion that specifically destroys the *edl* gene (Figure 3A, Table S2). Furthermore, we were able to rescue the cardiac phenotype of *edl* by introducing a genomic *edl* transgene ((Yamada et al., 2003); Figure 3E) and by artificially expressing *edl* in the dorsal mesoderm cells or in cardioblasts (Figure 3F,G), demonstrating that Edl is required directly within these cell types. In accordance, *edl* mRNA is found within the cardiogenic region during stages 10 to 11 and in CBs during stage 12 (Figure 3-figure supplement 1A-C). Thereafter *edl* expression shifts to the pericardial region, where it persists until stage 15 (Figure 3-figure supplement 1D and data not shown).

**Figure 3.**
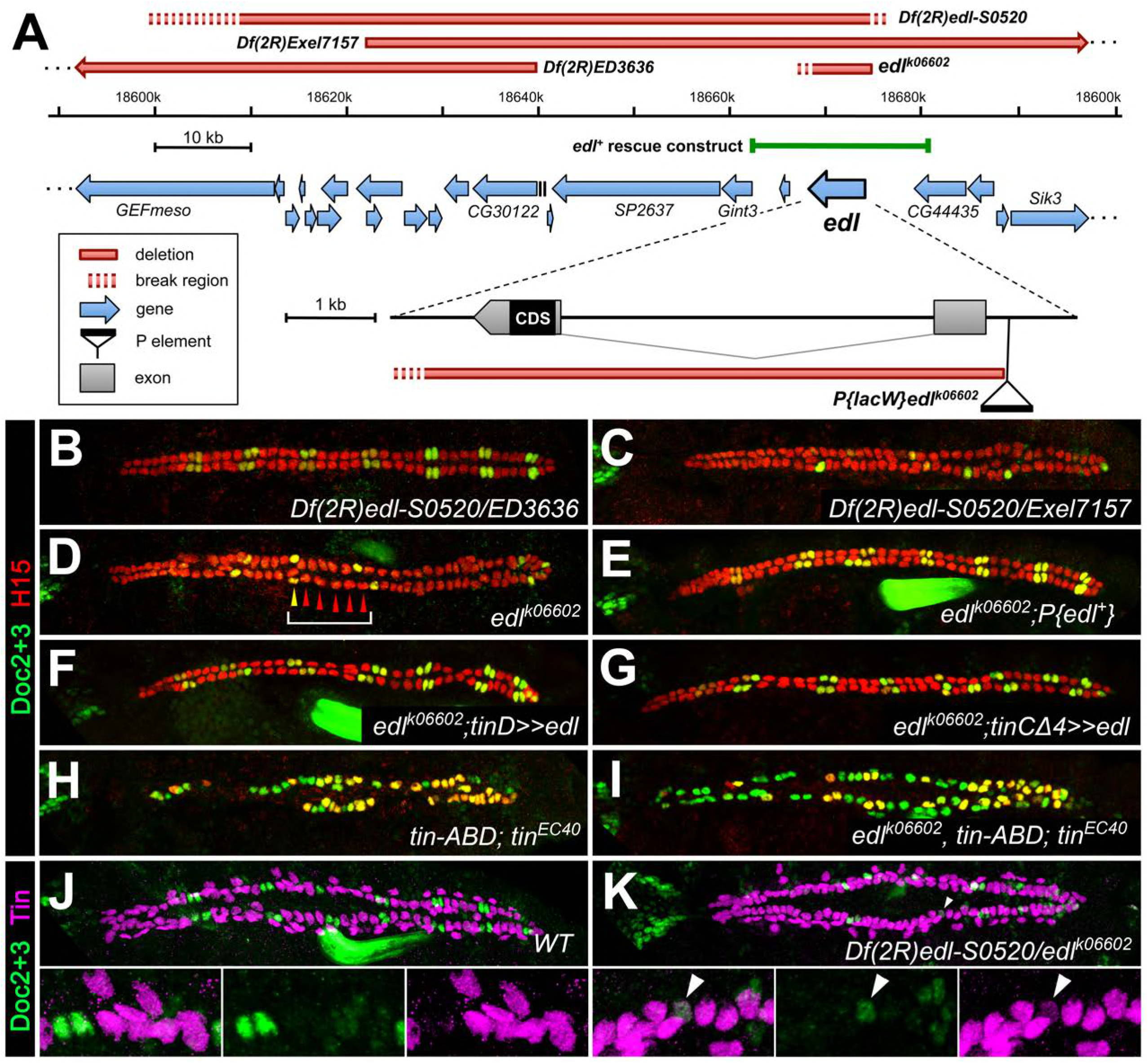
Edl is a decisive factor of ostial cardioblast specification. (A) Map of the *edl* locus with the used alleles and deficiencies. (B-I) Doc2+3/H15 stainings as in Figure 1. (B) Embryo with transheterozygous combination of *Df(2R)edl-S0520* (*edl*^*-*^) and *Df(2R)ED3636* (*edl*^*+*^) showing a regular "2+4" CB pattern of oCBs and gCBs. By contrast, amorphic *edl* mutants *Df(2R)edl-S0520/Exel7157* (C) and *edl*^*k06602*^ (D) have only few oCBs. Note the occurrence of "1+5" CB patterns (bracket). (E) The regular CB pattern is restored by a genomic *edl*^*+*^ transgene. A nearly normal CB pattern is observed in *edl* mutants upon expression of *UAS-edl* in the dorsal mesoderm via *tinD-GAL4* (F) or only in CBs or their progenitors via *tinCΔ4-GAL4* (G). In cardioblast-specific *tin* mutants (carrying a rescue construct for early *tin* function) all CBs present become Doc^+^, irrespective of whether *edl* is functional (H) or not (I). Observation of some H15^-^ Doc^+^ CBs in (H) and (I) suggest that robust H15 expression requires normal *tin* function. (J) Mutually exclusive expression of Doc and Tin proteins in the wild type at late stage 15. (K) In *edl* mutants, Doc and Tin are co-expressed at low levels in some CBs (arrowhead).

A distinctive feature of *edl* mutants is that the normal "2+4" pattern of 2 Doc^+^ CB + 4 Doc^-^ CBs is often transformed into a "1+5" pattern (e.g. bracket in Figure 3D), indicating a fate switch from ostial to generic CBs. However, Edl is not a direct activator of *Doc* expression because Doc is found in CBs of *edl* double mutants with CB-specific ablation of *tin* (Figure 3I), a phenotype reminiscent of that of CB-specific *tin* single mutants (Figure 3H; (Zaffran et al., 2006)). This suggests that *edl* normally contributes to the activation of *Doc* in oCBs via suppression of *tin*. This role of *edl* in CB patterning is further supported by the observation of some CBs with low levels of Tin and Doc in *edl* mutants (Figure 3K; compare to the strictly complementary distribution of Doc and Tin in the wild type, Figure 3J).

Next we analyzed Edl function by ectopic expression. Consistent with a mesoderm-autonomous function, overexpressing *edl* in the dorsal ectoderm via *pnr^MD237^-GAL4* has no significant effect on cardiogenesis (data not shown). By contrast, overexpression of *edl* in the entire mesoderm via *twist-GAL4* results in an increase of CB numbers (Figure 4A) and a decrease of OPCs (described in the next subsection). The increase in Doc^+^ CBs is disproportionately high. The extra Doc^+^ CBs in the heart proper also activate ostial cell differentiation markers such as *wg* (data not shown). In agreement with the proposed function of Edl as a negative regulator of PntP2 (Yamada et al., 2003), our overexpression phenotypes of *edl* are very reminiscent to that of *pntP2*-specific (*pnt*^*RR112*^ reported in (Alvarez et al., 2003) and *pnt*^*MI03880*^ shown in Figure 4B) and amorphic *pnt* mutants (*pnt*^*Δ88*^, *pnt*^*2*^; see Figure 4E,I and (Alvarez et al., 2003)). Accordingly, overexpression of constitutively active PntP2^VP16^ (Figure 4C) or PntP1 (not shown) via *tinD+tinCΔ4-GAL4* causes a phenotype similar to that of *edl* loss-of-function mutants (Figure 3C,D). By contrast, analogous overexpression of the potential Edl target Yan/Aop leads to a loss of heart cells irrespective of their subtype. These losses may result from a more general block in cell specification and differentiation since Yan has been related to such functions in several other types of MAPK-dependent progenitors (Bidet et al., 2003; Caviglia & Luschnig, 2013; Halfon et al., 2000; Rebay & Rubin, 1995). If the predominant function of Edl during CB specification is the inhibition of Pnt, *edl pnt* double mutants should mimic *pnt* mutants. In principle, this is what we observed (Figure 4E,F; quantifications in Figure 4I). By contrast, *edl aop* double mutants show an additive combination of *aop* and *edl* single mutant phenotypes (compare Figure 4H with 4G and 3D; see also quantifications in Figure 4I). Amorphic *aop* mutants display a reduction in CB number irrespective of CB subtype, which we ascribe to a permissive function during CB development that is probably linked to its well-documented role in restricting *eve* expression in the early dorsal mesoderm ((Bidet et al., 2003; Halfon et al., 2000; Liu et al., 2006; Webber, Zhang, Mitchell-Dick, & Rebay, 2013)). Importantly, and in contrast to *edl* and *pnt* activity changes, manipulating *aop* activities does not lead to significant shifts in the oCBs:gCBs ratio (Figure 4I). Thus we suggest that Edl acts mainly via negative modulation of PntP2 activity during cardioblast diversification.

**Figure 4.**
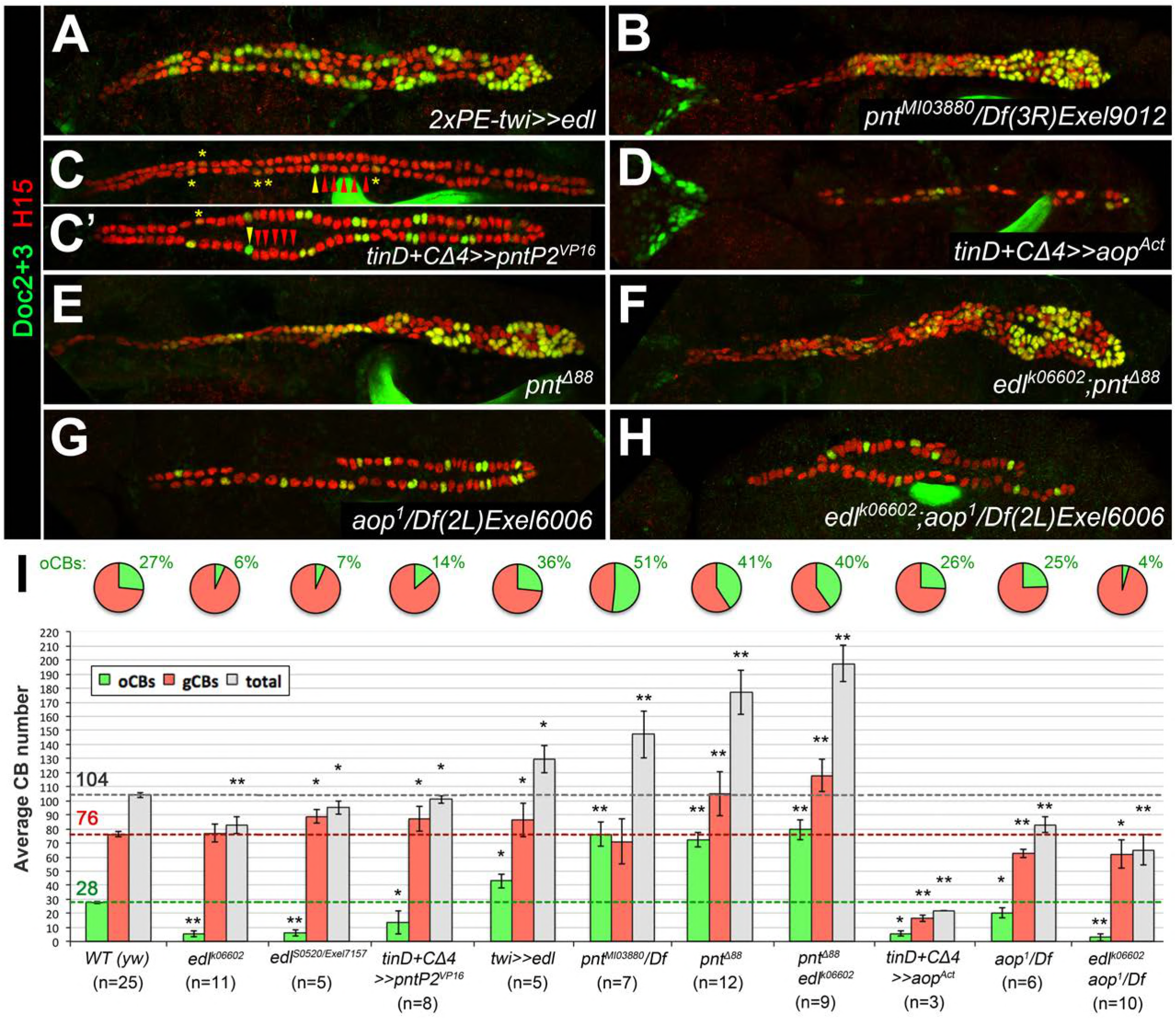
Edl promotes oCB fate via inhibition of PntP2. (A-H) CB pattern in embryos with modified activity of *edl* and/or genes encoding the ETS proteins Pnt and Yan revealed by H15/Doc2+3 stainings. (A) Pan-mesodermal *edl* overexpression via *twist-GAL4* leads to extra CBs with a disproportionately high increase in oCB numbers. This phenotype is reminiscent to that of the *pnt* mutants *pnt*^*MI03880*^ (a PntP2-specific mutant; here in trans with a *pnt*-deleting deficiency, B) and *pnt*^*Δ88*^ (without any functional Pnt isoform, E). (C,C') Conversely, an *edl* mutant-like phenotype (loss/conversion of oCBs, exemplified by arrowheads for one hemisegment, and CBs with low Doc levels marked by asterisks) is generated by overexpression of a constitutively active PntP2 variant in the dorsal/cardiogenic mesoderm. C and C' depict strong and weak phenotypes, respectively. (D) Overexpression of the constitutively active repressor Yan/Aop leads to a loss of gCBs and oCBs. (E,F) The CB phenotypes of *pnt* and *edl pnt* double mutants are very similar suggesting that *edl* acts mainly by blocking Pnt activity during CB specification. (G) Hemizygous *aop* mutant showing a moderate reduction of both CB types. (H) *edl aop* double mutant combining *aop*-like and *edl*-like defects. (I) Quantification of cardioblasts in various genotypes affecting Edl, Pnt or Yan/Aop activities (annotated as in Figure 1M).

### Edl and Pnt regulate ostial fate by controlling *seven-up* expression

The population of oCBs is characterized by expression of *svp*. In *svp* mutants all oCBs are converted into Tin^+^/Doc^-^ CBs due to de-repression of *tin* ((Gajewski et al., 2000; Lo & Frasch, 2001; Zaffran et al., 2006); Figure 5-figure supplement 1A). Therefore we tested the possibility that Edl promotes oCB fate by regulating *svp*. In the wild type, expression of *svp* is recapitulated by the enhancer trap *svp*^*AE127*^*-lacZ* (Figure 5A; Lo, 2001 #1072). In *edl* mutants, *svp*-LacZ expression is strongly reduced in cardiac cells (Figure 5B,D). The reduction in numbers of both *svp*-LacZ^+^ oCBs and OPCs at late stages (Figure 5D cf. 5C) suggests that *edl* already affects the fates of their common progenitors. Consistent with a function in promoting *svp* expression and oCBs fates, mesodermal overexpression of *edl* leads to larger numbers of *svp*-LacZ^+^ cardiac cells, particularly of CBs, where *svp* expression correlates with expanded Doc expression (Figure 5E,F). As shown for *Doc* expression, *svp* expression can be suppressed by PntP2 hyperactivity (green asterisks in Figure 5H). These observations and further evaluation of the epistatic relations between *svp* and *edl* (Figure 5-figure supplement 1) demonstrate that *edl* affects CB patterning by blocking Pnt activity upstream of *svp*.

**Figure 5.**
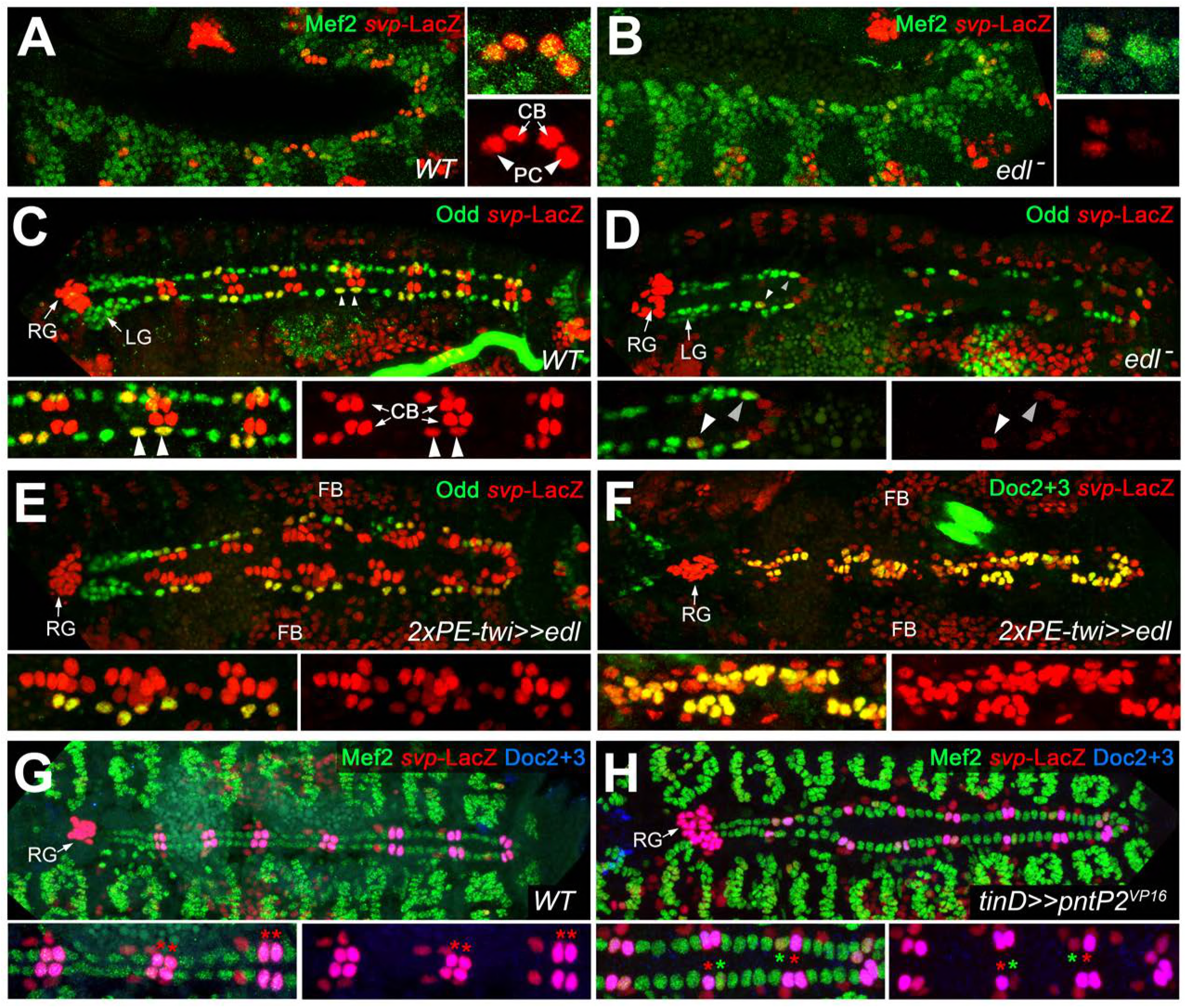
Edl is required for *svp* expression. (A) In stage 12 control embryos (lateral view) carrying one copy of *svp*^*AE127*^*-lacZ*, β-galactosidase is detected in oCBs (arrows) and their sibling OPCs (arrowheads) within the Mef2-labeled mesoderm. (B) Cardiac *svp-*LacZ expression is strongly reduced in *edl* mutants (*Df(2R)edl-S0520/Exel7157;svp*^*AE127*^*-lacZ/+*). (C-E) Odd/*svp*-LacZ staining in stage 16 embryos. (C) In the control, each hemisegment contains two oCB-related *svp*-LacZ^*+*^ OPCs and two *svp*-LacZ^*-*^ OPCs. The total number of OPCs decreases if *edl* is absent (*Df(2R)edl-S0520/edl-L19;svp*^*AE127*^*-lacZ/+*) (D) or overexpressed (E), but different OPC subpopulations account for these losses: *svp*-LacZ^+^ OPCs are reduced in *edl* mutants, *svp*-LacZ^*-*^ OPCs in *edl* overexpressing embryos. (E,F) Pan-mesodermal overexpression of *edl* leads to a drastic increase in the number of *svp*-LacZ^*+*^/Doc^+^ cardioblasts (Odd^-^). Compare F to the control in Figure 2E. (G,H) Mef2/Doc/β-galactosidase staining in *svp-lacZ/+* controls (G) and embryos overexpressing constitutively active *pntP2*^*VP16*^ in the dorsal mesoderm (H). Overexpression of *pntP2*^*VP16*^ leads to reduced levels of *svp* and *Doc* expression (examples labeled with green asterisks) as compared to normal oCBs (red asterisks).

### Cardioblast subtype-specific expression of the PntP1 isoform is regulated by PntP2 and Edl

Proposing a gCB-specific function of Pnt, we next analyzed its cardiac expression. Boisclair Lachance et al. previously reported that the expression of a fully functional genomic *pnt-GFP* transgene mirrors the combined expression of all Pnt isoforms (Boisclair Lachance et al., 2014). The authors detected Pnt-GFP fusion protein in nearly all cells of the cardiac region, but highest levels were observed in two Yan-negative clusters per hemisegment flanking Eve^+^ cells. We confirmed and refined these observations showing that high levels of Pnt-GFP are present in the nuclei of gCB progenitors as identified by their position, characteristically enlarged size, presence of only low levels of Doc, and absence of *svp-*LacZ expression (Figure 6A). We attribute these high total Pnt levels largely to a gCB-specific expression of the PntP1 isoform since PntP1-specific antibodies (Alvarez et al., 2003) specifically label gCB progenitors (Figure 6B), whereas *pntP2* transcripts are present in a rather uniform pattern in the mesoderm including the cardiogenic area ((Klämbt, 1993) and data not shown). We further speculated that PntP2 could activate *pntP1* transcription in gCB progenitors for a sustained signaling response as found in other tissues (Shwartz, Yogev, Schejter, & Shilo, 2013). This assumption is indeed supported by our genetic data. First, we detect PntP1 in an expanded pattern in the cardiogenic mesoderm of *edl* mutants in which PntP2 activity is assumed to increase (Figure 6C). Second, overexpression of *edl* (i.e. repression of PntP2 function) as well as genetic disruption of *pntP2* resulted in a near-complete loss of cardiac PntP1 (Figure 6D,E; note persistent expression of PntP1 in other cells located more laterally). We conclude that the combined activities of Edl and PntP2 lead to the confined *pntP1* expression in gCBs. The EGF Spitz appears to be a major, although not necessarily the sole factor for the MAPK-mediated activation of PntP2 in this context, because PntP1 levels are reduced but not eradicated in cardiac cells of amorphic *spi* mutants (Figure 6F).

**Figure 6.**
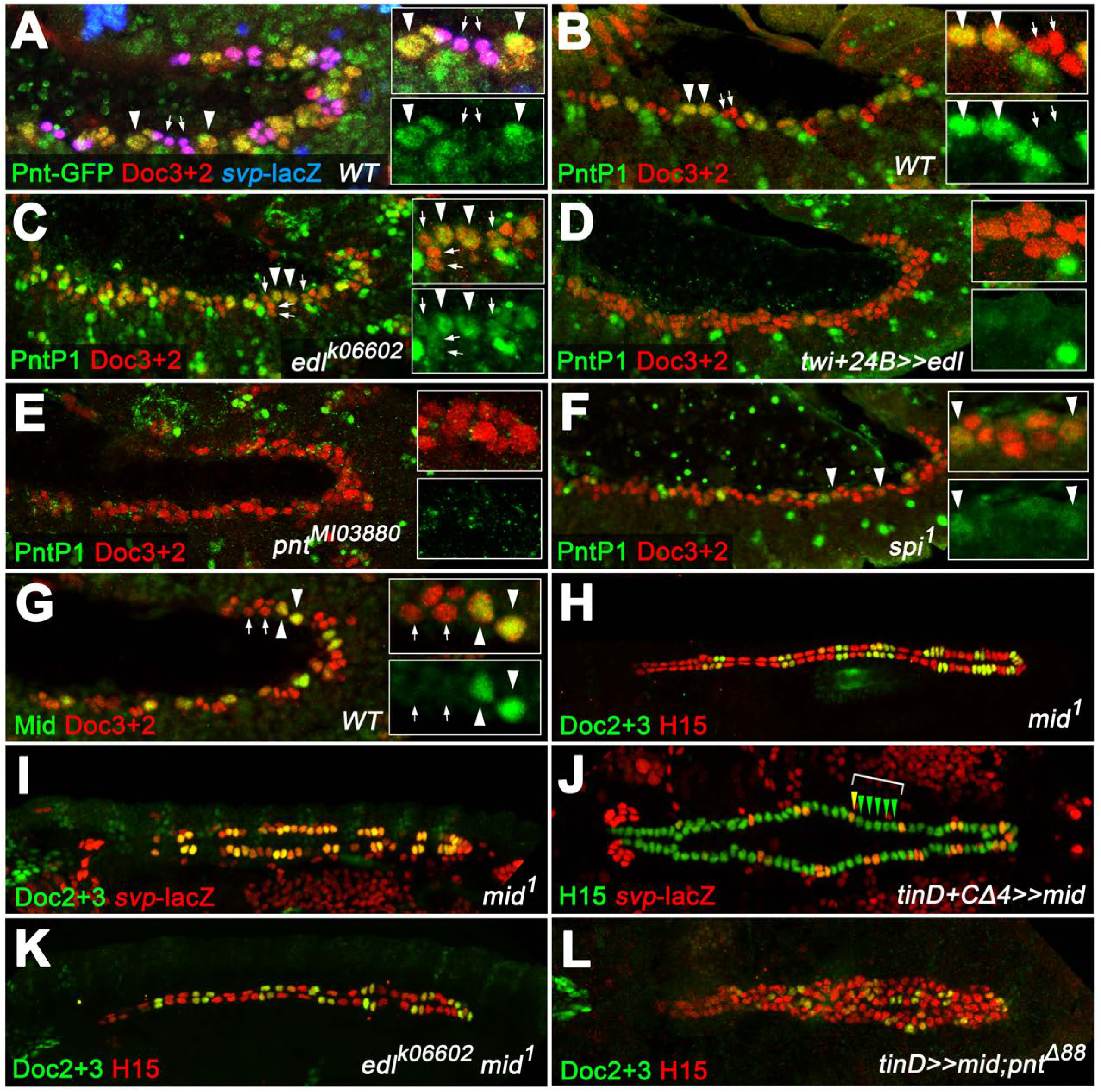
PntP1 and Mid are specifically expressed in early gCB progenitors to antagonize oCB fate. (A) Detection of Doc2+3, β-galactosidase and GFP-tagged Pnt (all isoforms) in a *pnt-GFP/+; svp-lacZ/+* embryo at the beginning of stage 12 (lateral view). Highest levels are observed in gCB progenitors (large *svp*-LacZ-negative nuclei with low levels of Doc, arrowheads) and low levels in oCBs and their siblings (small *svp*-LacZ^+^ nuclei with higher Doc levels, arrows). (B) At the onset of germ band retraction, PntP1 becomes expressed in gCB progenitors (arrowheads) of wild type embryos. Cardiac cells are labeled via anti-Doc3+2 staining. PntP1 is not detected in oCBs and their siblings (arrows). (C) In *edl*^*-*^ mutants cardiac PntP1 expression is generally increased and detected ectopically in some small nuclei that correspond to prospective oCBs and their siblings (arrows). (D) Pan-mesodermal overexpression of *edl* leads to a strong decrease of cardiac PntP1 expression while other mesodermal tissues are less affected. (E) The same effect is seen in *pntP2* mutants. (F) In *spi* mutants PntP1 levels are reduced as well, although not as severely as upon loss of *pntP2* function. (G) Like PntP1, Mid protein is found in gCB progenitors (arrowheads), but not in prospective oCBs (arrows) at the beginning of germ band retraction. (H,I) The cardiac phenotype of *mid* mutants is characterized by variable expansion of Doc, which largely correlates with ectopic *svp* expression in CBs (I, normal pattern shown in Figure 2E). (J) Overexpression of *mid* represses *svp* expression in H15-labeled cardioblasts (arrowheads indicate a hemisegment with five lacZ-negative nuclei). (K) Combining homozygous *mid* and *edl* mutations results in the restoration of oCBs in comparison to *edl* single mutants (Figure 3D), suggesting that *edl* normally antagonizes *mid* function. An additional *edl* function regarding the total CB number is not rescued by abrogation of *mid*. (L) Overexpression of *mid* in the dorsal mesoderm via *tinD-GAL4* in a *pnt* null background converts many of the extra oCBs into gCBs (cf. Figure 4E).

### The Tbx20 ortholog Midline contributes to Pnt-dependent repression of *svp* in the working myocardial lineage

According to the common view, we expect Pnt to act as a transcriptional activator also during CB diversification, particularly since overexpression of PntP2 fused to the VP16 activator domain has essentially the same effect on cardiac patterning as PntP1 overexpression (Figure 5H and data not shown). Therefore, its negative impact on *svp* expression is likely to involve Pnt-dependent activation of a transcriptional repressor. Interestingly, the T-box factor Midline (Mid), like PntP1, shows expression in early gCB progenitors (Figure 6G). We previously reported that *mid* functions to maintain *tin* expression in gCBs, thereby restricting *Doc* expression to oCBs (Reim et al., 2005). Consistent with this function our EMS screen also generated novel *mid* alleles showing the same CB patterning defects as previously described alleles (Table S1, Figure 6H and data not shown). While a direct regulation of *tin* by Mid was previously proposed to be responsible for these changes (supported by the gain- and loss-of-function phenotypes of *mid*; (Qian et al., 2005; Reim et al., 2005)), another non-exclusive scenario could involve repression of *svp* by Mid. Consistent with the latter, we observe a *Doc*-like expansion of *svp* expression in *mid* loss-of-function mutants (Figure 6I) and a reduction of *svp* expression upon ectopic overexpression of *mid* in cardiac cells (Figure 6J). The cardiac pattern phenotype of *edl mid* double mutants is a composite of the single mutant phenotypes. The number of oCBs (average oCBs: 24.4 ±3.6; n=6) is strongly increased as compared to *edl* mutants, but reduced in comparison with *mid* mutants, with total CB numbers being similar to those of *edl* mutants. In some cases a near wild type pattern is observed (Figure 6K), although many embryos display an asymmetric arrangement of CBs. While the prevalence of many Doc-negative CBs in this background implies that *mid* is not the only factor that limits oCB fate, it also indicates that *edl* is normally required in the oCB lineage to restrict *mid* activity, possibly by blocking a Pnt-dependent activation of *mid* transcription. This hypothesis is indeed supported by the reversion of ectopic *Doc* and *svp* expression in *pnt* mutants upon forced *mid* expression (Figure 6L, Figure 6-figure supplement 1C). By contrast, overexpression of the potential Mid target *tin* in this background only represses *Doc*, but not *svp* (Figure 6-figure supplement 1D).

To further test the idea that Mid is a repressor of oCB fate downstream of *pnt*, we analyzed whether it is a direct target of Pnt. Notably, an enhancer identified as a Tin target and named *midE19* (*mid180* for a shorter minimal version) was recently shown to drive *mid* expression specifically in gCBs ((Jin et al., 2013; Ryu, Najand, & Brook, 2011); Figure 7A). Consistent with our assumption that this enhancer is also a target of Pnt, very little *midE19*-GFP activity is detectable in *pnt* mutants (Figure 7B), reduced activity is observed in embryos with mesodermal *edl* overexpression (Figure 7-figure supplement 1A), and expanded activity is seen upon overexpression of PntP1 (Figure 7C; note occasional expansion into CBs with no detectable Tin) or PntP2^VP16^ (not shown). An observed reduction of *midE19*-driven GFP levels in many of the retained Tin^+^ gCBs of *rho* mutants (Figure 7-figure supplement 1B) corroborates that EGF signaling feeds into *mid* activation. The idea that *mid* is a target of Pnt is further supported by the almost complete elimination of reporter activity upon mutating a single ETS binding site within the mid180 minimal cardiac enhancer (Figure 7D,E) as well as the strong reduction of endogenous *mid* transcription in emerging CBs during germ band retraction stages in *pnt* mutants (Figure 7F-I). After germ band retraction, endogenous *mid* is activated independent of *pnt* in all CBs (Figure 7K) as observed in the wild type (Figure 7J) indicating that distinct mechanisms regulate *mid* transcription in early gCB progenitors and maturing CBs.

**Figure 7.**
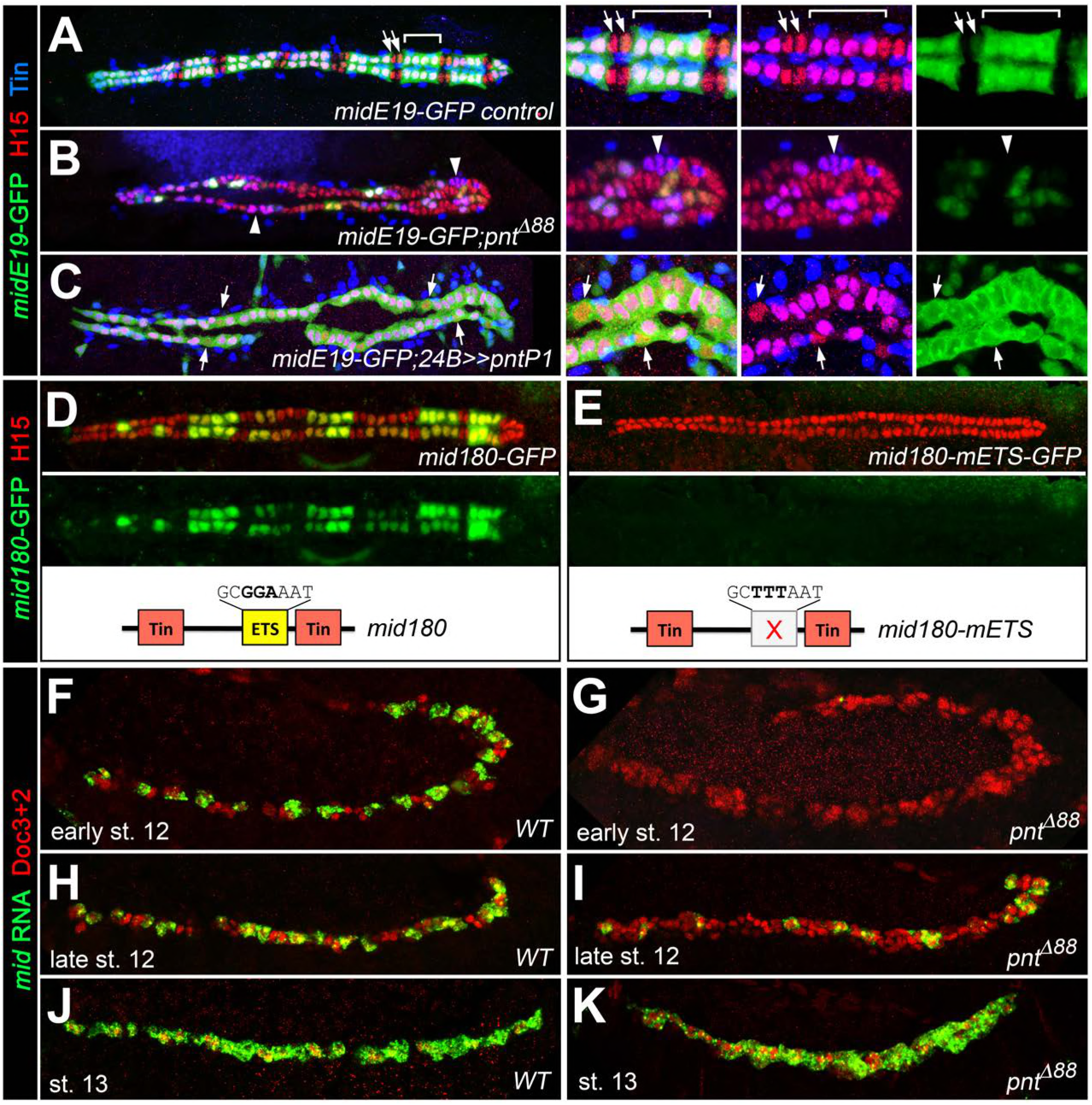
Characterization of a Pnt-responsive *mid* enhancer. (A-C) Stainings for GFP (green), H15 (red) and Tin (blue) in stage 16 embryos carrying the *midE19-GFP* reporter. (A) In a wild type heart, *midE19*-GFP is strongly expressed in the four pairs of Tin^+^/H15^+^ gCBs (bracket), whereas no or very little GFP is detectable in Tin^-^/H15^+^ oCBs (arrows). (B) Despite an overall increase in CB number, *midE19*-GFP expression is severely reduced in amorphic *pnt* mutants. Most of the Tin^+^/H15^+^ gCBs (purple nuclei, arrowheads) lack GFP expression. (C) Overexpression of *pntP1* via *how*^*24B*^*-GAL4* leads to nearly continuous *midE19*-GFP expression in CBs. In some instances the reporter is activated even in Tin^-^ CBs (arrows). (D) Expression of GFP driven by the minimal cardiac *mid* enhancer, *mid180*, is less robust than *midE19*- GFP but shows essentially the same expression pattern. The minimal enhancer contains a single ETS binding motif flanked by two Tin binding sites (indicated in the scheme below). (E) Mutating the ETS binding site leads to near-complete abolishment of *mid180*-GFP expression. (F-K) Analysis of *mid* mRNA expression in cardiac cells doubly stained with anti-Doc3+2 antibody. In the wild type, *mid* mRNA is first detected in gCB progenitors at early stage 12 (F); its expression begins to expand during germ band retraction (H) until it reaches continuous expression in all CBs at stage 13 (J). By contrast, amorphic *pnt* mutants show reduced cardiac *mid* expression during germ band retraction (G,I). Regular uniform *mid* expression is observed only after germ band retraction (K).

In sum our data lead to the conclusion that EGF signaling contributes to gCB specification by at least two distinct mechanisms, Pnt-independent specification of a subset of cardiac progenitors as well as Pnt-dependent inhibition of ostial cardioblast fate. Modulation by Edl is needed to inhibit Pnt-dependent gene activation and thus enable formation of ostial cardioblasts.

## Discussion

The specification and diversification of particular cell types are linked to the establishment of lineage-specific transcriptional programs. The differences in these programs are often prompted by distinct local signaling activities. The cells in the early heart fields of *Drosophila* acquire their cardiogenic potential by intersecting BMP and Wnt signal activities (Frasch, 1995; Reim & Frasch, 2005; X. Wu et al., 1995), but cell diversification within this area requires additional regulatory inputs. Previous studies established that progenitors of cardioblasts, pericardial cells and dorsal somatic muscles are selected by RTK/Ras/MAPK signaling, whereas lateral inhibition by Delta/Notch signaling activity counteracts this selection in neighboring non-progenitor cells (Carmena et al., 2002; Grigorian et al., 2011; Hartenstein et al., 1992). The progenitors of the definitive cardiogenic mesoderm, which give rise to all cardiac cells except for the somatic muscle lineage-related EPCs, co-express the cardiogenic factors Tin, Doc and Pnr, a unique feature that separates them from other cells (Reim & Frasch, 2005). In addition to limiting the number of progenitors, Notch signaling has a second function during *Drosophila* cardiogenesis that promotes pericardial (or in thoracic segments, hematopoietic) over myocardial fate (Albrecht, Wang, Holz, Bergter, & Paululat, 2006; Grigorian et al., 2011; Hartenstein et al., 1992; Mandal, Banerjee, & Hartenstein, 2004). Other factors previously reported to impose heterogeneity in the heart field include the cross-repressive activities of the homeodomain factors Eve and Lbe (T. Jagla, Bidet, Da Ponte, Dastugue, & Jagla, 2002) as well as ectoderm-derived Hedgehog (Hh) signals (Liu et al., 2006; Ponzielli et al., 2002). In segmental subsets of cardioblasts, Hh signaling was proposed to act as a potential activator of *svp* in prospective oCBs (Ponzielli et al., 2002) but whether these are direct or indirect effects of Hh on these cells has not been ascertained.

Based on the findings of our study, we present a novel model of cardioblast diversification that introduces EGF signaling activities and lineage-specific modulation of the MAPK effector Pointed by Edl as crucial factors for the specification of generic working myocardial and ostial cell fates.

### A novel model for cardioblast diversification connecting EGF signaling, ETS protein activity and lineage-specific transcription factor patterns

Combining previous findings with our new data we have conceived the regulatory model of cardioblast diversification illustrated in Figure 8. We propose that EGF/MAPK signaling promotes the development of generic working myocardial progenitors (red/left cell in Figure 8) by two mechanisms that differ in their requirement for the ETS protein Pnt:

1) EGF promotes the correct selection and specification of gCB progenitors. This is evident from our loss- and gain-of-function analysis of EGF signaling components. Notably, in most hemisegments of the analyzed EGF pathway mutants the number of gCBs is reduced by even numbers, indicating a defect prior to completion of the final mitotic division at the progenitor stage. This EGF function is obviously independent of *pnt*, since *pnt* null mutants display excessive numbers of CBs (with gCB numbers comparable to the wild type or even increased), a phenotype different from that of mutants defective in EGF pathway components upstream of Pnt ((Alvarez et al., 2003) and this study).

2) EGF signals affect the diversification of CB progenitors by impinging on a PntP2-dependent transcriptional cascade that eventually leads to suppression of Tin^-^ oCB and the adoption of Tin^+^ gCB fates. This function is mediated by stimulating the gCB progenitor-specific expression of regulatory genes such as *mid* (depicted in red in Figure 8), which in turn will promote transcription of gCB-specific differentiation genes and/or repression of oCB-specific factors (depicted in green in Figure 8).

**Figure 8.**
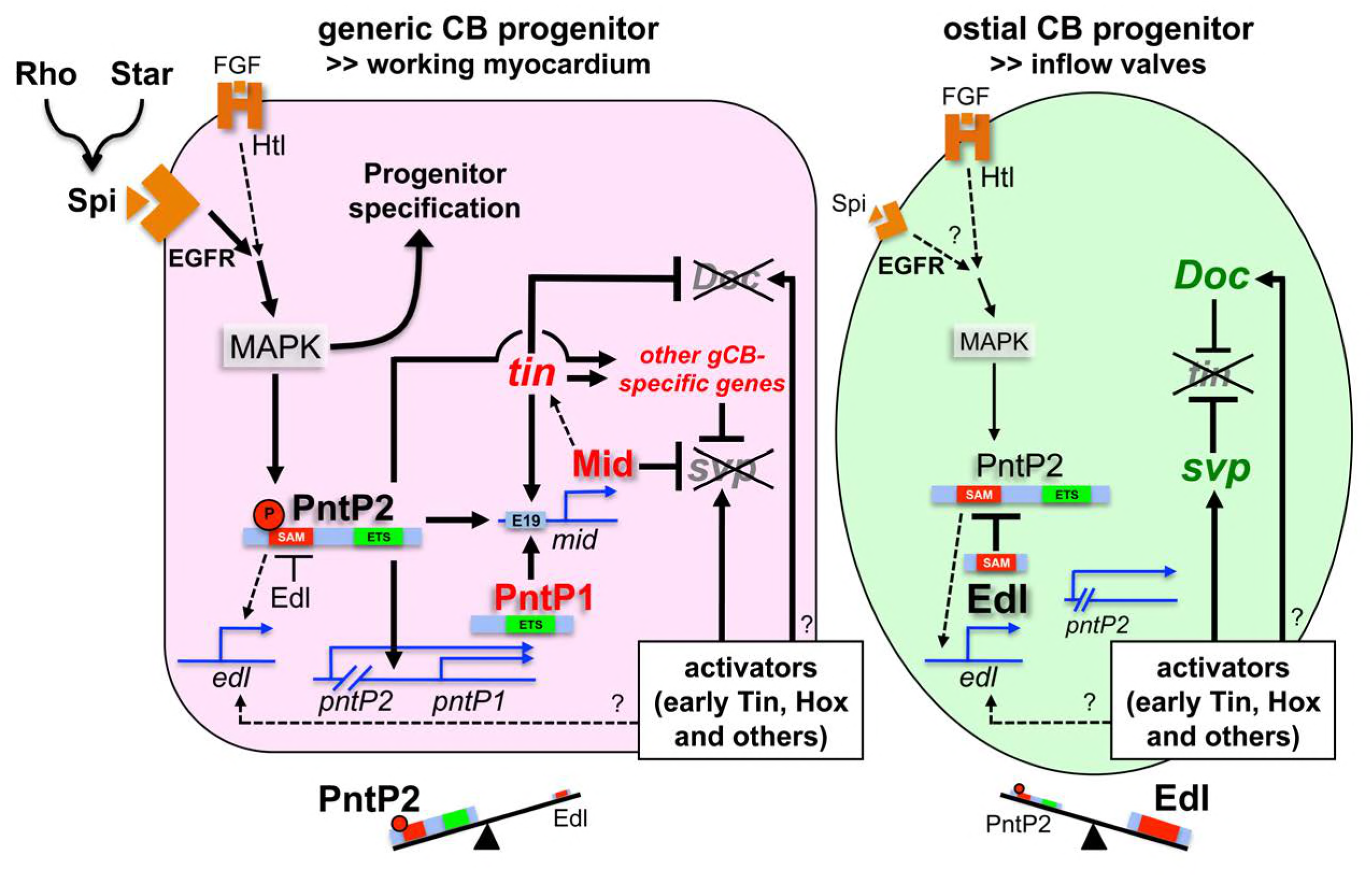
Model of regulatory interactions in generic and ostial CB progenitors. Genes activated in a subtype-specific manner in gCB or oCB progenitors are colored in red and green, respectively. Larger font sizes and thicker lines indicate higher levels. Dashed lines indicate presumed regulations. In principle, MAPK can be activated in cardiac progenitors by EGF/EGFR and FGF/Htl signals. Generic cardioblast development depends on EGF-activated MAPK signaling which provides *pnt*-independent and *pnt*-dependent functions. The suppression of *svp* and subsequent regulation of *tin* and *Doc* is a *pnt*-dependent function that is in part mediated by direct activation of *mid* via the midE19 enhancer in presumptive gCBs. This step is likely to be supported by the gCB-specific expression of constitutive active PntP1. The gCB-specific cascade may require a higher level of MAPK activity to overcome the blockage of PntP2 by Edl. Alternatively or in addition, Edl levels might be differentially regulated in gCBs and oCBs by yet unknown mechanisms. In oCB progenitors, Edl keeps activated PntP2 below a critical threshold leading to absence or delayed onset of expression of oCB fate antagonists such as *mid*. This in turn permits *svp* activation by Hox genes and Tin derived from early stages. Presumed transcriptional activators of *svp* acting downstream of segmental Hh signals in oCB progenitors are not mandatory in this model, although it does not categorically exclude such contributions.

#### Basic features of gene regulation in the gCB lineage

We identified *mid* as a key target gene of Pnt in gCB progenitors based on its early gCB-specific expression, Pnt-dependent transcriptional regulation and its ability to repress the oCB-specific regulator gene *svp*. Since Svp represses *tin* expression (Gajewski et al., 2000; Lo & Frasch, 2001), *svp* suppression provides an important part of the explanation for the previously reported positive role of Mid in maintaining *tin* expression in gCBs (Qian et al., 2005; Reim et al., 2005), although it does not exclude the possibility that Mid stimulates *tin* expression also directly. Of note, the vertebrate Mid ortholog Tbx20 is also a promoter of working myocardial fate that can act as transcriptional activator and repressor depending on context (Cai et al., 2005; M. K. Singh et al., 2005; Stennard et al., 2003; Stennard et al., 2005; Takeuchi et al., 2005). While Tin acts as a repressor of *Doc* via unknown mechanisms in gCBs, it does not repress *svp* ((Zaffran et al., 2006) and Figure 6-figure supplement 1D). On the contrary, at least in the early cardiogenic mesoderm, it acts as an activator of *svp* in oCB progenitors (Ryan, Hendren, Helander, & Cripps, 2007). Thus, in the absence of appropriate repressors such as Mid, *svp* expression can expand into gCBs.

#### Basic features of gene regulation in the oCB lineage

In prospective oCB progenitors, Pnt activity must be kept in check to permit *svp* expression and thereby *tin* repression and *Doc* activity. Fittingly, we identified *edl*, a gene linked to negative regulation of MAPK signaling and cell identity determination in several tissues - including the eye (Yamada et al., 2003) and recently in certain somatic muscle progenitors (Dubois, Frendo, Chanut-Delalande, Crozatier, & Vincent, 2016) - as a novel regulator in the context of cardiac cell specification, particularly that of oCB progenitor fate (green/right cell in Figure 8). This function was first hinted at by the over-proportional increase of *svp*-expressing oCBs in *pnt* mutants (Alvarez et al., 2003). Our phenotypic analysis demonstrates that Edl is required for *svp* and *Doc* gene activity (the latter being due to restriction of *tin* expression) as well as the restriction of PntP2-dependent PntP1 expression in cardiac progenitors. Molecularly, Edl can modulate the activities of PntP2 as well as Yan (Baker et al., 2001; Qiao et al., 2006; Qiao et al., 2004; Tootle et al., 2003; Vivekanand et al., 2004; Yamada et al., 2003). The comparison of single and double mutant phenotypes, combined with the reproducibility of nearly all aspects of the cardiac *pnt* phenotype by Edl overexpression, implies that Edl acts primarily by inhibiting Pnt during cardiac cell diversification, although we cannot fully exclude additional interactions with Yan. Our observations further support the function of Edl as an antagonist of Pnt (first demonstrated in the context of eye and chordotonal organ development, (Yamada et al., 2003)) and rule out an initially proposed Pnt-stimulating function (Baker et al., 2001).

#### Linkage of MAPK and Pnt activities

The involvement of Edl also leads to important conclusions regarding the placement of Pnt function within the cardiac gene regulatory network. Based on the phenotypic discrepancies between *pnt* and other EGF pathway components (gain and loss of CBs, respectively), Alvarez et al. proposed that PntP2 acts independent of MAPK signaling to limit the number of CBs (Alvarez et al., 2003). Since we found that Edl blocks Pnt activity in oCB progenitors, and Edl is thought to antagonize PntP2 mainly by blocking MAPK-dependent phosphorylation (Qiao et al., 2006), we propose that PntP2 acts downstream of MAPK also during cardiogenesis (see Figure 8). This is further supported by our data demonstrating *spi*-sensitive cardiac expression of PntP1 and the observation that, if timed properly, both EGF and Pnt activities can lead to expanded gCB and reduced oCBs populations. However, not all MAPK activities require *pnt*, which is the case for the pro-cardiogenic activities of EGF. Notably, parallel *pnt*-dependent and *pnt*-independent MAPK signaling functions take place also during other processes such as epithelial branching morphogenesis (Cabernard & Affolter, 2005).

### EGF signaling and cardiac progenitor selection

As discussed above, cardioblast formation as such is independent of *pnt*. How could this be achieved? Growth factor-activated MAPK can also phosphorylate the repressor Yan thereby diminishing its activity as an antagonist of progenitor selection (Halfon et al., 2000; O'Neill et al., 1994; Rebay & Rubin, 1995). Therefore, it is conceivable that MAPK activity in the context of CB progenitor selection might be primarily required to eliminate the repressive activity of Yan. This would be consistent with the observed reduction of cardiac cells upon *aop/yan* hyperactivation ((Halfon et al., 2000) and this study). In this context, a minor function of Edl could contribute to the robustness of cardiac progenitor selection and thus total cardioblast and pericardial cell numbers by reducing the repressive Yan activity.

According to our data, EGF signals are the major source for MAPK activation and progenitor specification in the symmetrically dividing progenitors of gCBs and OPCs. By contrast, EGF signals are dispensable (in high doses even unfavorable) for the development of progenitors of oCBs and their sibling OPCs. Thus, EGF signaling appears to be more critical for cell fate specification of cardiac cells than for their mere survival. Our overexpression studies demonstrate that the timing of EGF signals is crucial for this function. In previous studies, earlier functions of MAPK signaling might have obscured its specific impact on gCBs and OPC subtypes. While early pan-mesodermal activation of MAPK signaling or expression of constitutive active Pnt forms via the *twi-GAL4* driver reduces the numbers of all cardiac cells except the Eve^+^ progenitors ((Alvarez et al., 2003; Bidet et al., 2003; Liu et al., 2006) and our own data), later MAPK activation favors formation of the symmetrically dividing OPC and gCB progenitor subpopulations (based on our experiments with *tinD-GAL4-*driven *rho*; increased total numbers of OPCs and CBs were also observed after expressing activated forms of Ras or the receptors of FGF and EGF with a *Mef2-GAL4* driver, (Grigorian et al., 2011)). We propose that the specification of these progenitors requires the context of the definitive cardiogenic mesoderm, whereas premature MAPK activation in all mesoderm cells negates any pro-cardiogenic effects due to the massive expansion of Eve^+^ clusters (which are normally the first cells in the heart field to display MAPK and *rho* activity) at the expense of the cardiac progenitors in the neighboring C14/C16 clusters (Buff et al., 1998; T. Jagla et al., 2002; Liu et al., 2006; Qian et al., 2005); and our own data not shown).

### Special features of Pnt-dependent regulation in working myocardial cells

Our model of CB diversification incorporates the observation that the PntP1 isoform is activated specifically in gCB progenitors in a PntP2-dependent and EGF-sensitive fashion. This is reminiscent of the situation in other tissues such as the developing eye where the PntP1 isoform is also activated in a MAPK/PntP2-dependent manner (Gabay et al., 1996; O'Neill et al., 1994; Shwartz et al., 2013). We propose that PntP1 becomes activated at a particular threshold of MAPK/PntP2 activity. This activation marks a point of no return for CB diversification, because PntP1 cannot be inhibited via Edl. The activation of PntP1 also explains why *edl* overexpression with relatively late acting drivers such as *tinD-GAL4* (as used in the *edl* mutant rescue experiment) does not cause the cardiac phenotypes observed with early pan-mesodermal drivers.

Besides *pntP1* and the already discussed *mid* gene, there are very likely additional target genes activated by PntP2 and/or PntP1 to execute the differentiation program in generic working myocardial cells. Incomplete conversion of gCBs in *mid* mutants also calls for the existence of additional repressors that contribute to oCB fate suppression. Interestingly, a study investigating Tin target genes found that cardiac target enhancers of Tin are not only enriched for Tin binding sites but also for a motif highly reminiscent of ETS binding sites, termed "cardiac enhancer enriched (CEE) motif" (with the consensus ATT[TG]CC or GG[CA]AAT in antisense orientation) (Jin et al., 2013). Mutation of four CEE sites (one of which overlapping our predicted ETS binding site) in a ca. 600 bp version of the *midE19* enhancer nearly abolished reporter activity in that study. Thus, many of the CEE-containing Tin target enhancers might in fact also be targets of Pnt (potentially mediating ETS-dependent activation) or Yan (potentially mediating ETS-dependent repression in the absence of MAPK signals). Therefore, a combination of closely spaced Tin and ETS binding sites might be a key signature in enhancers of working myocardial genes, although additional features must be present in their architecture to distinguish them from Tin+ETS-binding site containing enhancers active in pericardial cells or their progenitors (Halfon et al., 2000). The differences might include elements directly or indirectly regulated by Delta/Notch signaling. Notably, the juxtacrine Notch ligand Delta is upregulated in the CB lineage in an MAPK activity-dependent manner (Grigorian et al., 2011). Hence, it is conceivable that Pnt proteins might stimulate *Delta* transcription in gCBs to control OPC development in a non-autonomous manner. This would explain both, simultaneous mis-specification of gCB progenitors and non-ostial-related OPCs in EGF mutants as well as phenotypic similarities between *pnt* mutants and mutants for components of the Delta-Notch signaling pathway. However, Notch pathway mutants do not display a biased increase of oCBs ((Albrecht et al., 2006) and our own data not shown) because of the herein described function of Pnt in suppressing *svp* transcription and oCB fate.

### What is the original signal that discriminates generic and ostial progenitors?

Our model proposes that factors that tilt the balance between PntP2 activity and Edl will have a major impact on CB subtype choice (see Figure 8). Thus any input that modestly increases MAPK/PntP2 activity within the appropriate window of time would favor gCB fate, whereas factors that have the opposite effect should promote oCB specification. This points to activities that impinge on the highly complex and dynamic expression of *rho*, because the Rhomboid protease is a key determinant in the decision of which cells will activate the more broadly expressed EGF Spitz and thus emanate signaling activity. A candidate would be Hh, which was proposed to be an oCB-promoting factor (Ponzielli et al., 2002). However, its effect on MAPK and *rho* activity in the dorsal mesoderm was suggested to be positive rather than negative (Liu et al., 2006). This would refute a function favoring oCB fate. The regulation of *rho* and the role of *hh* during CB diversification await more detailed analysis in future studies.

Factors that regulate *edl* expression levels might also determine the outcome of the competition between Edl and Pnt. The *edl* gene was found to be positively regulated by EGF signaling, and a target of Pnt and Yan, and thus was proposed to provide a negative feedback system for EGF inputs (Baker et al., 2001; Leatherbarrow & Halfon, 2009; Vivekanand et al., 2004; Yamada et al., 2003). Our model therefore includes regulation by Pnt as a possibility (dashed arrows in Figure 8). ChIP-on-chip experiments suggest that *edl* is also targeted by cardiogenic factors (Junion et al., 2012).

The spatio-temporal dynamics and detailed mechanisms that regulate MAPK and *edl* activities within the cardiogenic mesoderm remain to be investigated in future studies. Such studies may also help to understand lineage decisions in other tissues and species. Edl/Mae-relatives are also present in non-Dipteran insects (e.g. *Tribolium*, (Bucher & Klingler, 2005)), echinoderms, and the chordate *Ciona*. Although no clear ortholog of Edl appears to be present in vertebrates, a SAM domain-only isoform of the human Yan-relative TEL2 as well as *Drosophila* Edl were shown to inhibit transcriptional stimulation by the mammalian Pnt orthologs ETS1/ETS2 in cell culture (Gu et al., 2001; Vivekanand & Rebay, 2012). Hence, the restriction of ETS protein activities by protein-protein interactions offers an intriguing mechanism to fine-tune MAPK signaling output in developing tissues of both invertebrates and vertebrates.

## Materials and Methods

### *Drosophila melanogaster* stocks

The mutants *Df(2R)edl-S0520, Egfr*^*S0167*^, *Egfr*^*S2145*^, *Egfr*^*S2307*^, *Egfr*^*S2561*^, *mid*^*S0021*^, *mid*^*S2961*^, *numb*^*S1342*^, *numb*^*S3992*^, *numb*^*S4439*^, *pyr*^*S3547*^ (Reim, Hollfelder, Ismat, & Frasch, 2012), *spi*^*S3384*^, *Star*^*S4550*^ were recovered from our EMS screen. The lines *mid^1^, UAS-mid-B2, how^24B^-GAL4, pnr^MD237^-GAL4, svp^AE127^-lacZ* (a *svp* mutant in homozygous condition), *UAS-svp.I, 2xPE-twi-GAL4, twi-SG24-GAL4, tinD-GAL4, UAS-tin#2, {tin-ABD}T003-1B1; tin^EC40^, UAS-p35* were as described previously (Reim et al., 2012; Reim et al., 2005; Zaffran et al., 2006). In addition the following strains were used: *aop*^*1*^*=aop*^*IP*^ (Nüsslein-Volhard, Wieschaus, & Kluding, 1984; Rogge et al., 1995), *UAS-aop.ACT-IIa* (Rebay & Rubin, 1995), *edl^L19^=Df(2R)edl-L19* (*edl* and some neighboring genes deleted) and *UAS-edl-X* (both from Y. Hiromi, (Yamada et al., 2003)), *P{lacW}edl*^*k06602*^ (Baker et al., 2001; Török, Tick, Alvarado, & Kiss, 1993), *UAS-Egfr^DN^.B-29-77-1;UAS-Egfr^DN^.B-29-8-1* (Buff et al., 1998), *htl*^*YY262*^ (Gisselbrecht, Skeath, Doe, & Michelson, 1996), *midE19-GFP* ((Jin et al., 2013); from M. Frasch), *pnt*^*Δ88*^ (Scholz, Deatrick, Klaes, & Klämbt, 1993), *pnt*^*MI03880*^ (PntP2-specific; harbors a gene-trap cassette with an artificial splice acceptor followed by stop codons upstream of the *pntP1* transcription start site, (Venken et al., 2011)), *UAS-pntP2^VP16^-2* ((Halfon et al., 2000); originally from C. Klämbt), *UAS-pntP1-3* and *UAS-pntP2-2* (Klaes, Menne, Stollewerk, Scholz, & Klämbt, 1994), *PBac{pnt-GFP.FPTB}VK00037* (R. Spokony and K. White, (Boisclair Lachance et al., 2014)), *pyr*^*18*^ and *ths*^*759*^ (Klingseisen, Clark, Gryzik, & Müller, 2009), *rho*^*7M43*^ (Jürgens, Wieschaus, Nüsslein-Volhard, & Kluding, 1984), *rho*^*L68*^ (Salzberg et al., 1994), *rho*^*EP3704*^ (Bidet et al., 2003), *UAS-rho(ve.dC)* (de Celis, Bray, & Garcia-Bellido, 1997), *spi*^*1*^=*spi*^*IIA14*^ (Nüsslein-Volhard et al., 1984), *Star*^*B0453*^ ((Chen et al., 2008), from F. Schnorrer), *tinCΔ4-GAL4* ((Lo & Frasch, 2001), from M. Frasch), *Df(2R)Exel7157*, and about 180 additional deficiencies spanning chromosome 2 (except where noted, all stocks available from the Bloomington Stock Center).

Flies expressing *edl*^*+*^ from a transgene were generated anew by standard P-element transgenesis using the previously described rescue construct *edl[+t18]* ((Yamada et al., 2003), provided by Y. Hiromi).

Unless noted otherwise, *y w* or *S-18a-13b-16c.1* control (Hollfelder et al., 2014) flies were used as wild type controls. Mutant lines were maintained over *GFP-* or *lacZ-*containing balancer chromosomes to allow recognition of homozygous embryos. Flies were raised at 25°C, except for UAS/GAL4-driven overexpression at 29°C.

### Isolation and mapping of novel EMS mutants

Novel EMS-induced mutants were obtained from our screen for embryonic heart and muscle defects and mapped to a particular gene through extensive complementation testing analogous to the previously described procedure (Hollfelder et al., 2014). Many alleles were mapped by unbiased complementation tests with a set of chromosome 2 deficiencies and subsequent non-complementation of lethality and embryonic phenotype by previously described alleles. *Df(2R)edl-S0520* was mapped by non-complementation of lethality with *Df(2R)Exel7157, Df(2R)edl-L19* and *Df(2R)ED3636*, but the cardiac phenotype was only reproduced *in trans* with *Df(2R)Exel7157, Df(2R)edl-L19* and *edl*^*k06602*^. Novel alleles of *Egfr* and *Star* were mapped using a candidate gene approach.

### Molecular analysis of mutations and deletions

Several EMS alleles and the unmutagenized *S-18a-13b-16c.1* control were analyzed by sequencing of overlapping PCR products covering the coding sequence and splicing sites of the candidate gene as described (Hollfelder et al., 2014). Details about the mutations are provided in Table S1. The area deleted by *Df(2R)edl-S0520* and its approximate break points were determined by iterative PCR amplification tests. The insertion of *P{lacW}edl*^*k06602*^ near the *edl* transcription start site was confirmed by PCR using primers binding to the 5' *P* end and adjacent genomic DNA. Although the integrity of the both *P* element ends could be confirmed by PCR, no genomic *edl* sequences expected next to the 3' *P* end could be amplified using several primer pairs shown to amplify control DNA. This indicates that *P{lacW}edl*^*k06602*^ is associated with a deletion in *edl*. Details of the deletion mapping are listed in Table S2.

### Generation of reporter constructs for enhancer analysis

The *mid180-GFP* reporter constructs were generated according to a similar *lacZ* construct published by Ryu et al. (Ryu et al., 2011). The forward primer 5'-*Eco*RI-CGTGCCTCCCACTTCAGGGCGG-3' and the backward primer 5'-*Bam*HI-TTAATTTCATTTTTCACTCTGCTCACTTGAGATTCCCCTGCTTTGTCTGCGGC<u>ATT***TCC***GC</u>TTCT-3' were used to amply DNA from *y w* flies. The predicted ETS binding site matching the antisense sequence of published ETS binding motifs ((Halfon et al., 2000; Hollenhorst, McIntosh, & Graves, 2011), underlined) was mutated by replacing the invariable *TCC* core (bold) with *AAA* in the backward primer. Amplicons were cloned into *Eco*RI/*Bam*HI of pH-Stinger-attB (Jin et al., 2013), sequenced and inserted into the *attP2* landing site via *nos*-driven ΦC31 integrase.

### Staining procedures

Embryo fixations, immunostainings for proteins and RNA in situ hybridizations were carried out essentially as described (Knirr, Azpiazu, & Frasch, 1999; Reim & Frasch, 2005). VectaStain Elite ABC kit (Vector Laboratories) and tyramide signal amplification (TSA, PerkinElmer Inc.) were used for detection of RNA and certain antigens (as indicated). The following antibodies were used: guinea pig anti-Doc2+3 (1:2000, TSA) and anti-Doc3+2 (1:1000) (Reim et al., 2003), rabbit anti-H15/Nmr1 (1:2000), guinea pig anti-H15/Nmr1 (1:2000), rabbit anti-Mid/Nmr1 (early stages: 1:250, TSA; late stages: 1:1000 direct) and rabbit anti-PntP1 (1:250, TSA) (all from J. Skeath; (Alvarez et al., 2003; Leal, Qian, Lacin, Bodmer, & Skeath, 2009)), rabbit anti-Mef2 (1:1500) (from H.T. Nguyen), rat anti-Odd (1:600, TSA) (Kosman, Small, & Reinitz, 1998), rabbit anti-Eve (1:3000) (Frasch, Hoey, Rushlow, Doyle, & Levine, 1987), rabbit anti-Tin (1:750) (Yin, Xu, & Frasch, 1997) (all from M. Frasch), rabbit anti-β-galactosidase (Promega, 1:1500), rabbit anti-GFP (Molecular Probes, 1:2000 and Rockland, 1:1000), mouse anti-GFP 3E6 (Life Technologies, 1:100, TSA), anti-cleaved-Caspase-3 (Asp175, Cell Signaling Technology, 1:100, TSA), sheep anti-Digoxigenin (Roche, 1:1000, TSA), monoclonal mouse antibodies anti-β-galactosidase 40-1a (1:20 direct or 1:50 with TSA), anti-Seven-up 5B11 (1:20, TSA) and anti-Wg 4D4 (1:30, TSA) (all from Developmental Studies Hybridoma Bank, University of Iowa), fluorescent secondary antibodies (1:200) (Jackson ImmunoResearch Laboratories and Abcam), biotinylated secondary antibodies (1:500) and HRP-conjugated anti-rabbit IgG (1:1000) (Vector Laboratories). TUNEL staining was performed as described (Reim et al., 2003) using the Millipore ApopTag S7100 kit in combination with TSA.

Digoxigenin-labeled antisense riboprobes against *mid, edl, rho* and *pntP2* were used for whole mount *in situ* hybridizations. The *mid* probe was generated as described previously (Reim et al., 2005). T7 promoter-tagged *edl, rho* and *pntP2* (isoform-specific exons) templates for *in vitro* transcription were generated by PCR (primers *edl*: CAATCGTGAAAGAGCGAGGGTC, T7-TGACGAGCAGAACTAAGGACTAGGC, *pnt*: CCAGCAGCCACCTCAATTCGGTC, T7-GCGTGCGTCTCGTTGGGGTAATTG, *rho*: ATGGAGAACTTAACGCAGAATGTAAACG, T7-TTAGGACACTCCCAGGTCG) from DNA of wild-type flies or flies carrying *UAS-rho(ve.dC*) or *UAS-pntP2*, respectively.

Embryos were mounted in Vectashield (Vector Laboratories). Images were acquired on a Leica SP5 II confocal laser scanning microscope and projected using Leica LAS-AF and ImageJ.

## Acknowledgements

We are grateful to Manfred Frasch for critical reading of the manuscript, Patrick Lo and Christoph Schaub for their contributions to the EMS screen, Edmar Heyland, Emi Vargatoth, Tanja Drechsler and Angela Bruns for technical assistance. We thank Manfred Frasch, Yasushi Hiromi, James Skeath, Hanh Nguyen, Frank Schnorrer, the Bloomington Stock Center and the Developmental Studies Hybridoma Bank (University of Iowa) for providing fly stocks or reagents.

## Author Contributions

BS, IR conceived, designed and performed the experiments, analyzed and interpreted the data, wrote the manuscript. DH, KS, LH performed the experiments, analyzed and interpreted the data.

## Supplementary information

**Supplementary Table 1.**
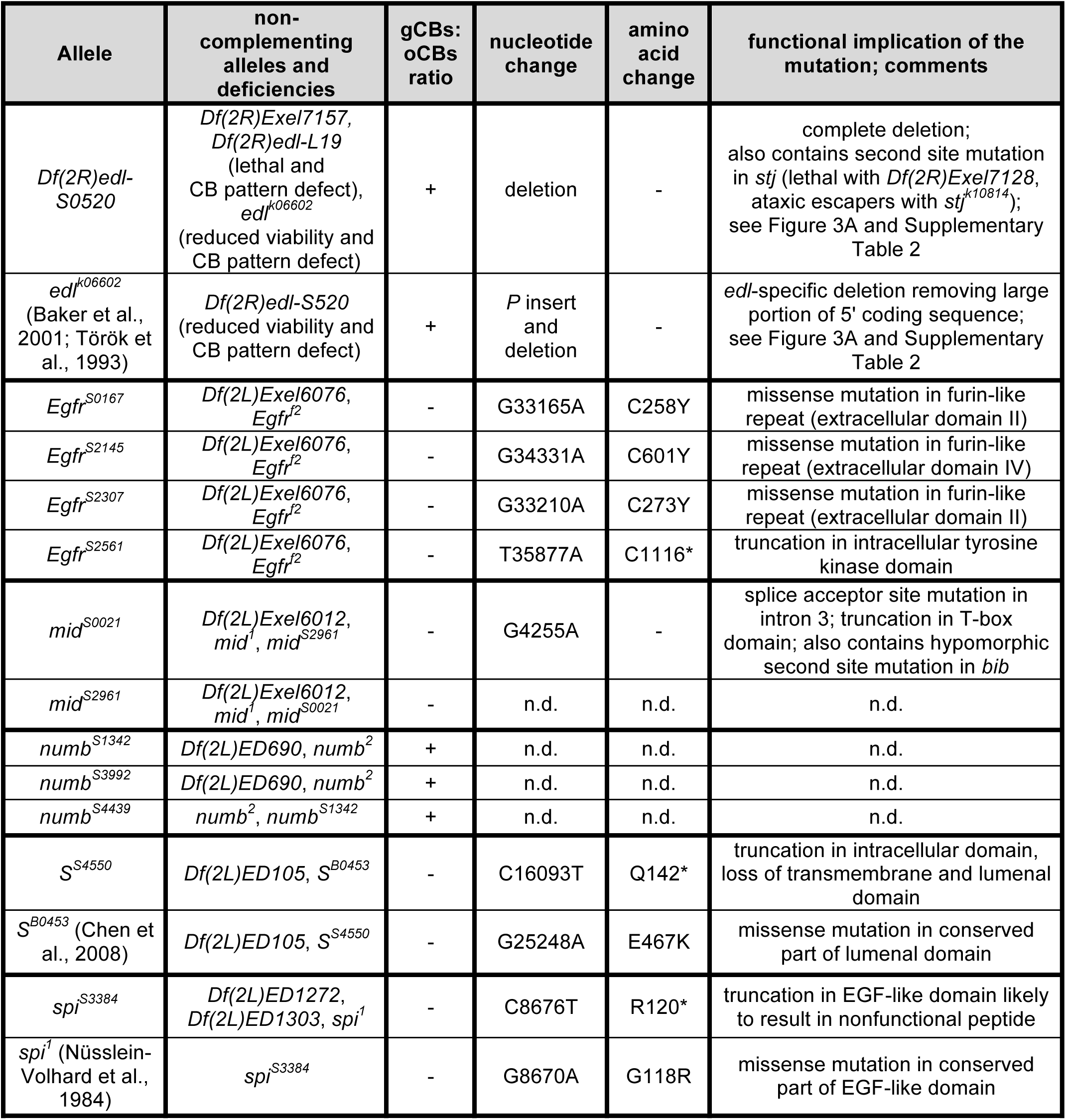
Alleles with cardioblast patterning defects isolated and/or characterized in this study. Unless indicated otherwise nucleotide positions are relative to transcription start site of transcript RA and amino acid positions of protein isoform PA (FB2017_01, released February 14, 2017; D. melanogaster R6.14); * indicates a nonsense mutation, n.d.: not determined.

| Allele | non-complementing alleles and deficiencies | gCBs: oCBs ratio | nucleotide change | amino acid change | functional implication of the mutation; comments |
| --- | --- | --- | --- | --- | --- |
| *Df(2R)edl-S0520* | *Df(2R)Exel7157, Df(2R)edl-L19* (lethal and CB pattern defect), *edl*<sup>k06602</sup> (reduced viability and CB pattern defect) | + | deletion | - | complete deletion; also contains second site mutation in *stj* (lethal with *Df(2R)Exel7128*, ataxic escapers with *stj*<sup>k10814</sup>); see Figure 3A and Supplementary Table 2 |
| *edl*<sup>k06602</sup> (Baker et al., 2001; Török et al., 1993) | *Df(2R)edl-S520* (reduced viability and CB pattern defect) | + | *P* insert and deletion | - | *edl*-specific deletion removing large portion of 5' coding sequence; see Figure 3A and Supplementary Table 2 |
| *Egfr*<sup>S0167</sup> | *Df(2L)Exel6076, Egfr*<sup>f2</sup> | - | G33165A | C258Y | missense mutation in furin-like repeat (extracellular domain II) |
| *Egfr*<sup>S2145</sup> | *Df(2L)Exel6076, Egfr*<sup>f2</sup> | - | G34331A | C601Y | missense mutation in furin-like repeat (extracellular domain IV) |
| *Egfr*<sup>S2307</sup> | *Df(2L)Exel6076, Egfr*<sup>f2</sup> | - | G33210A | C273Y | missense mutation in furin-like repeat (extracellular domain II) |
| *Egfr*<sup>S2561</sup> | *Df(2L)Exel6076, Egfr*<sup>f2</sup> | - | T35877A | C1116* | truncation in intracellular tyrosine kinase domain |
| *mid*<sup>S0021</sup> | *Df(2L)Exel6012, mid*<sup>1</sup>, *mid*<sup>S2961</sup> | - | G4255A | - | splice acceptor site mutation in intron 3; truncation in T-box domain; also contains hypomorphic second site mutation in *bib* |
| *mid*<sup>S2961</sup> | *Df(2L)Exel6012, mid*<sup>1</sup>, *mid*<sup>S0021</sup> | - | n.d. | n.d. | n.d. |
| *numb*<sup>S1342</sup> | *Df(2L)ED690, numb*<sup>2</sup> | + | n.d. | n.d. | n.d. |
| *numb*<sup>S3992</sup> | *Df(2L)ED690, numb*<sup>2</sup> | + | n.d. | n.d. | n.d. |
| *numb*<sup>S4439</sup> | *numb*<sup>2</sup>, *numb*<sup>S1342</sup> | + | n.d. | n.d. | n.d. |
| *S*<sup>S4550</sup> | *Df(2L)ED105, S*<sup>B0453</sup> | - | C16093T | Q142* | truncation in intracellular domain, loss of transmembrane and luminal domain |
| *S*<sup>B0453</sup> (Chen et al., 2008) | *Df(2L)ED105, S*<sup>S4550</sup> | - | G25248A | E467K | missense mutation in conserved part of luminal domain |
| *spi*<sup>S3384</sup> | *Df(2L)ED1272, Df(2L)ED1303, spi*<sup>1</sup> | - | C8676T | R120* | truncation in EGF-like domain likely to result in nonfunctional peptide |
| *spi*<sup>1</sup> (Nüsslein-Volhard et al., 1984) | *spi*<sup>S3384</sup> | - | G8670A | G118R | missense mutation in conserved part of EGF-like domain |

**Supplementary Table 2.**
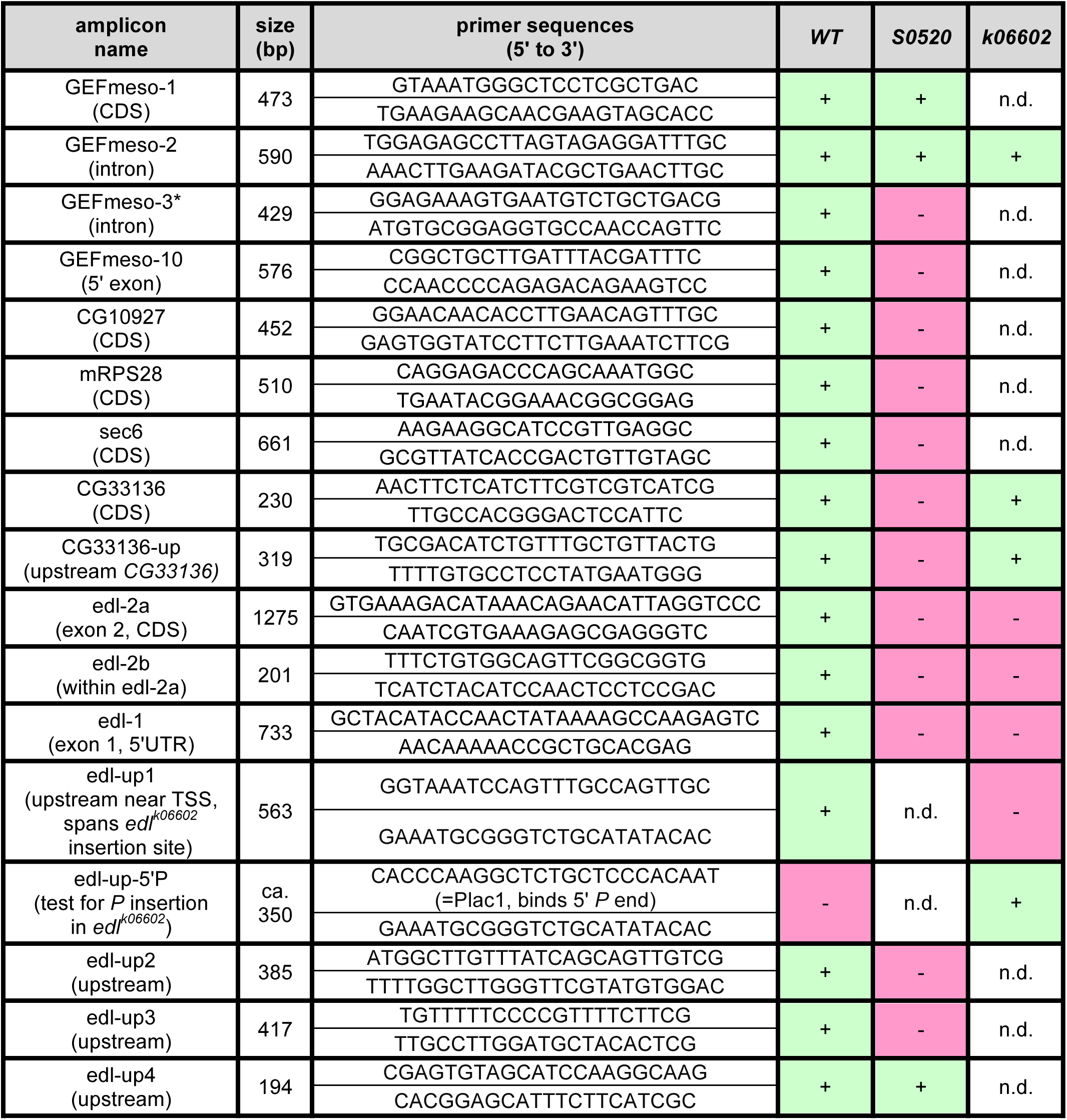
Characterization of *edl* deletions via PCR. Presence (+) or absence (-) of DNA fragments after PCR reaction including genomic DNA from homozygous *S-18a-13b-16c.1* control (*WT*), *Df(2R)edl-S0520* or *edl*^*k06602*^ animals and primer pairs as indicated. CDS: part of coding sequence, TSS: transcription start site, n.d.: not determined. Amplicons are listed in linear order as located on chromosome 2R. * Six additional intronic *GEFmeso* amplicons were also negative in *S0520*.

**Figure 1 - source data 1. Quantification of Doc^+^ oCBs, Doc^−^ gCBs and total cardioblasts.** data file: fig1-data1.xlsx

**Figure 1 - Figure supplement 1.**
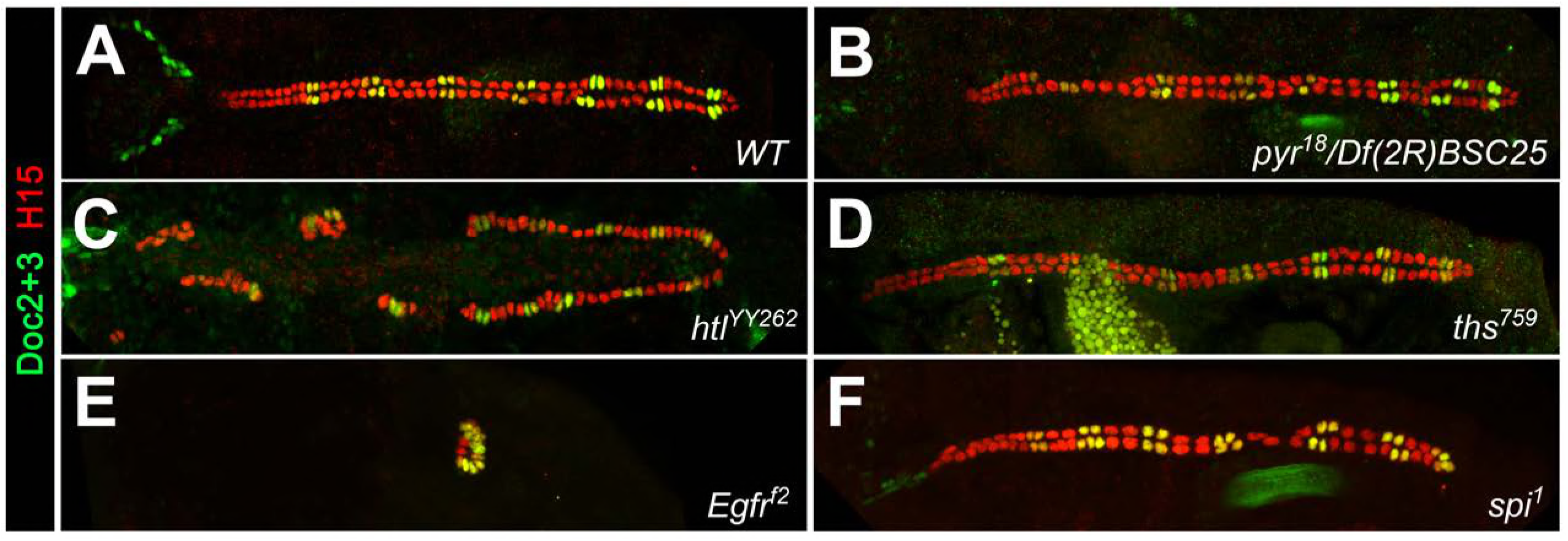
Cardiac patterning phenotypes in additional alleles of FGF and EGF pathway mutants. Embryos stained for H15 and Doc as in Figure 1. (A) Wild type with normal CB pattern. Reduced FGF/Htl signaling in *pyr^18^/Df(2R)BSC25* (*pyr*^*-/-*^ *ths*^*+/-*^) embryos (B) or homozygous mutants with the hypomorphic allele *htl*^*YY262*^ (C) leads to a random loss of gCBs and oCBs. (D) Neither significant changes in CB number nor patterning defects were observed in homozygous *ths*^*759*^ mutants. The mutants *Egfr*^*f2*^ (E) and *spi*^*1*^ (F) show essentially the same phenotypes as the corresponding alleles of the same genes shown in Figure 1.

**Figure 1 - Figure supplement 2.**
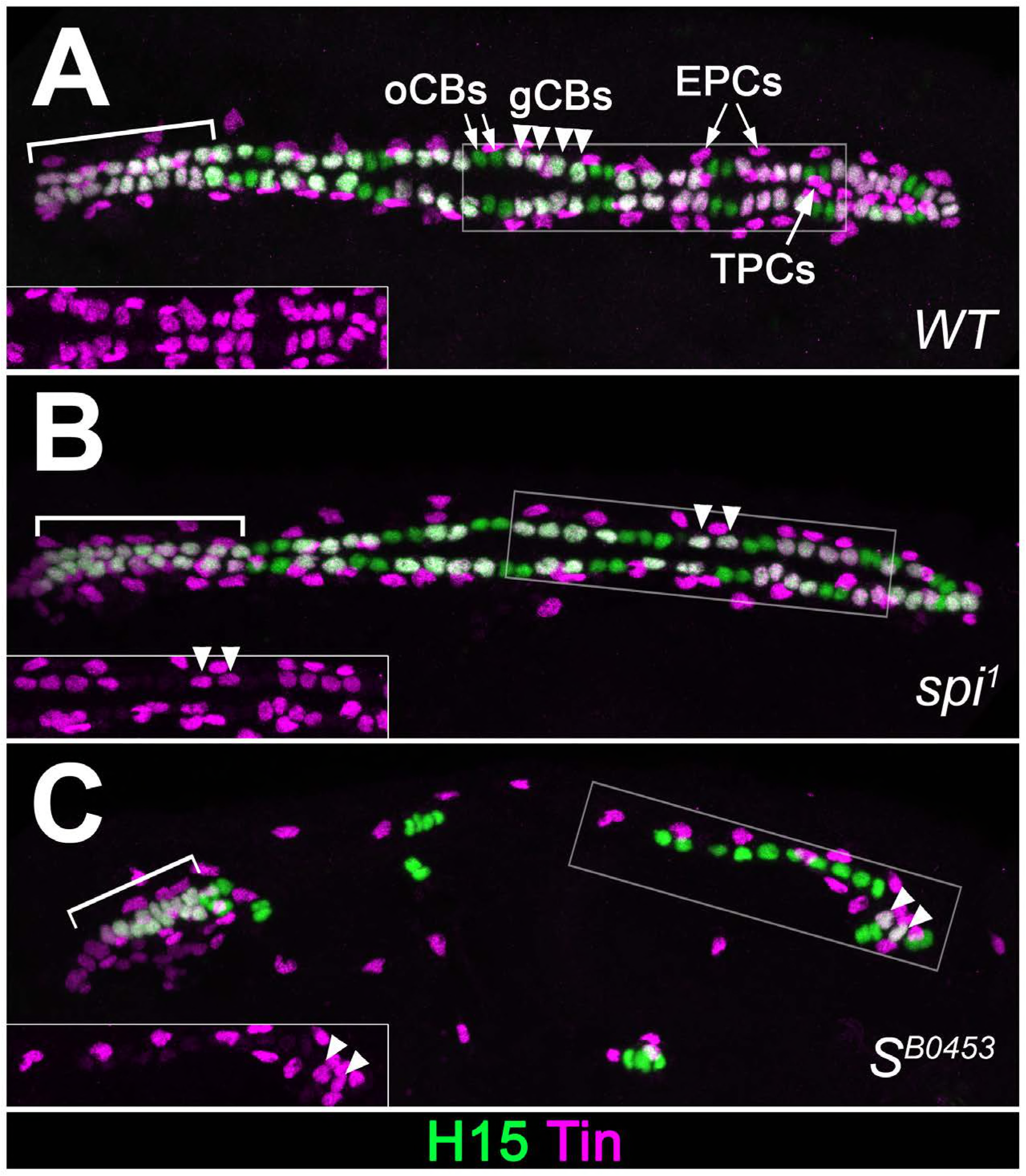
Extended analysis of cardiac patterning in EGF pathway mutants. Expression of Tin and H15 detected by immunostaining in stage 16 embryos. (A) In the wild type, each abdominal hemisegment contains four Tin^+^H15^+^ gCBs (white, arrowheads) and several Tin^+^H15^-^ PCs. In the anterior aorta (bracket) all CBs express Tin. (B) Homozygous *spi*^*1*^ mutant with reduced number of gCBs. (C) Homozygous *S*^*B0453*^ mutant in which only one pair of Tin^+^ gCBs has developed in abdominal segments. CBs of the anterior aorta (aa) are less affected. Correlating with the presence of Doc^+^ CBs, Tin^-^ CBs are present at near wild type numbers in *spi* and *S* mutants.

**Figure 1 - Figure supplement 3.**
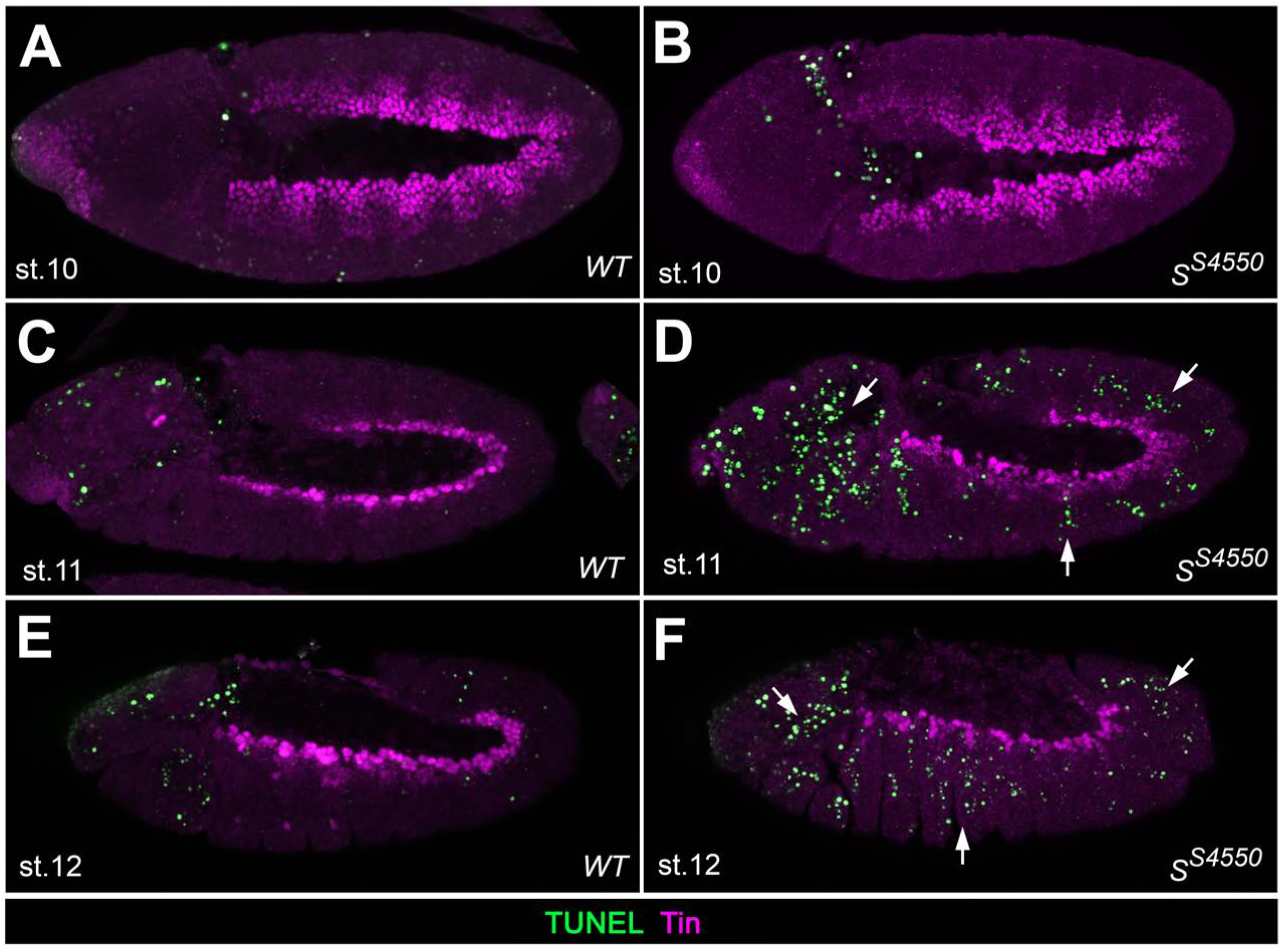
Analysis of apoptosis in *Star* mutants. TUNEL assay co-stained for Tin to detect apoptotic cells in the cardiogenic mesoderm of wild type embryos (A,C,E) and in amorphic *S* mutants (B,D,F) at the indicated stages. TUNEL signals are not found within in the Tin^+^ cardiogenic mesoderm of stage 10, 11 and 12 embryos in both wild type and mutant embryos, although such signals could readily detected in more ventral and lateral regions as well as in the head (arrows). Note the higher abundance of TUNEL staining in Tin-negative tissues in the mutants. This suggests that EGF signaling does not serve as a mere survival cue in the cardiogenic mesoderm but has a major function in specifying cardiac fates.

**Figure 1 - Figure supplement 4.**
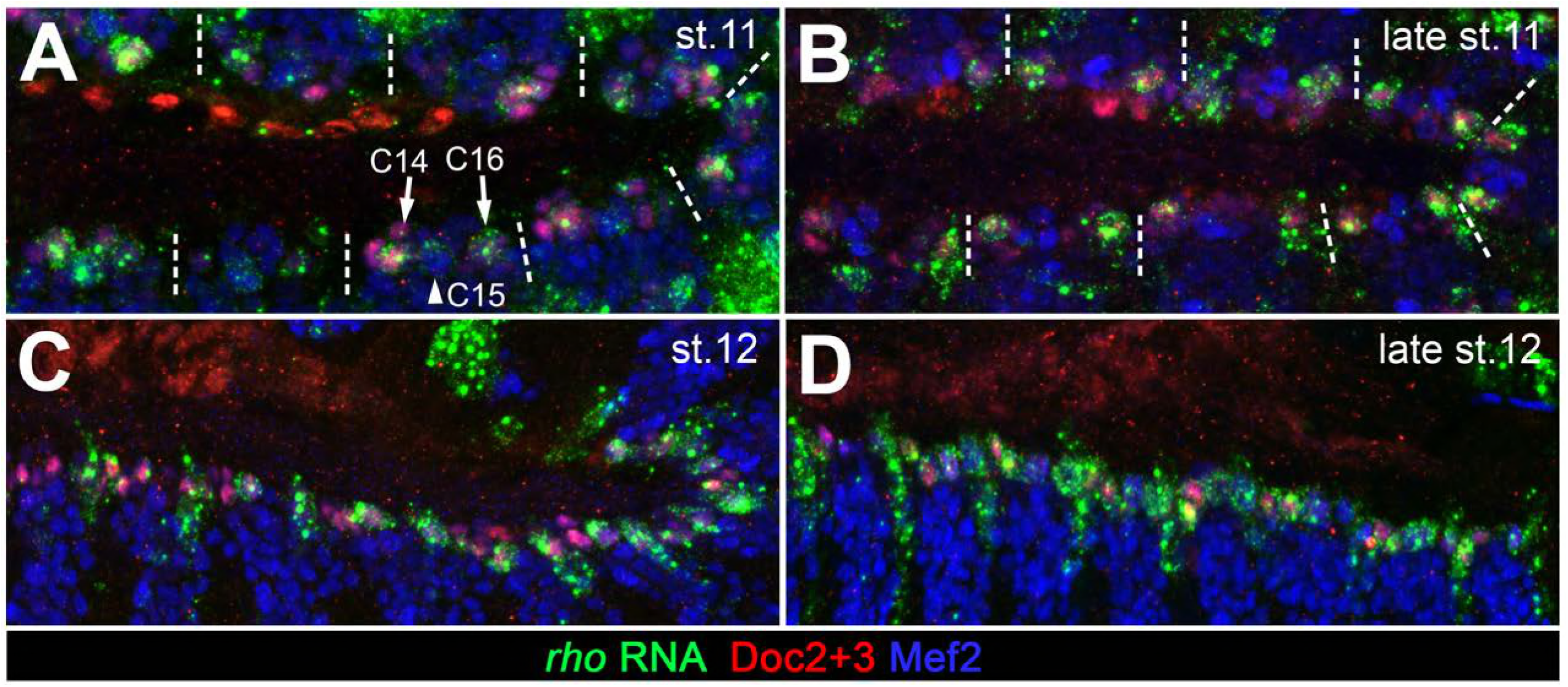
Expression of *rho* in cardiac cells. Detection of *rho* mRNA (green), Mef2 (blue) and Doc (red). (A) At stage 11, *rho* is detectable in clusters C14/C16 of the cardiac mesoderm and is fading from the central Doc-negative region containing EPC and somatic muscle progenitors. Some cardiac cells express higher levels than others. (B) At late stage 11, *rho* is expressed at high levels in one or a few cardiac progenitors close to the dorsal segment borders. (C, D) As cardioblasts align near the dorsal mesoderm margin during stage 12, *rho* continues to be expressed in most CBs.

**Figure 3 - Figure supplement 1.**
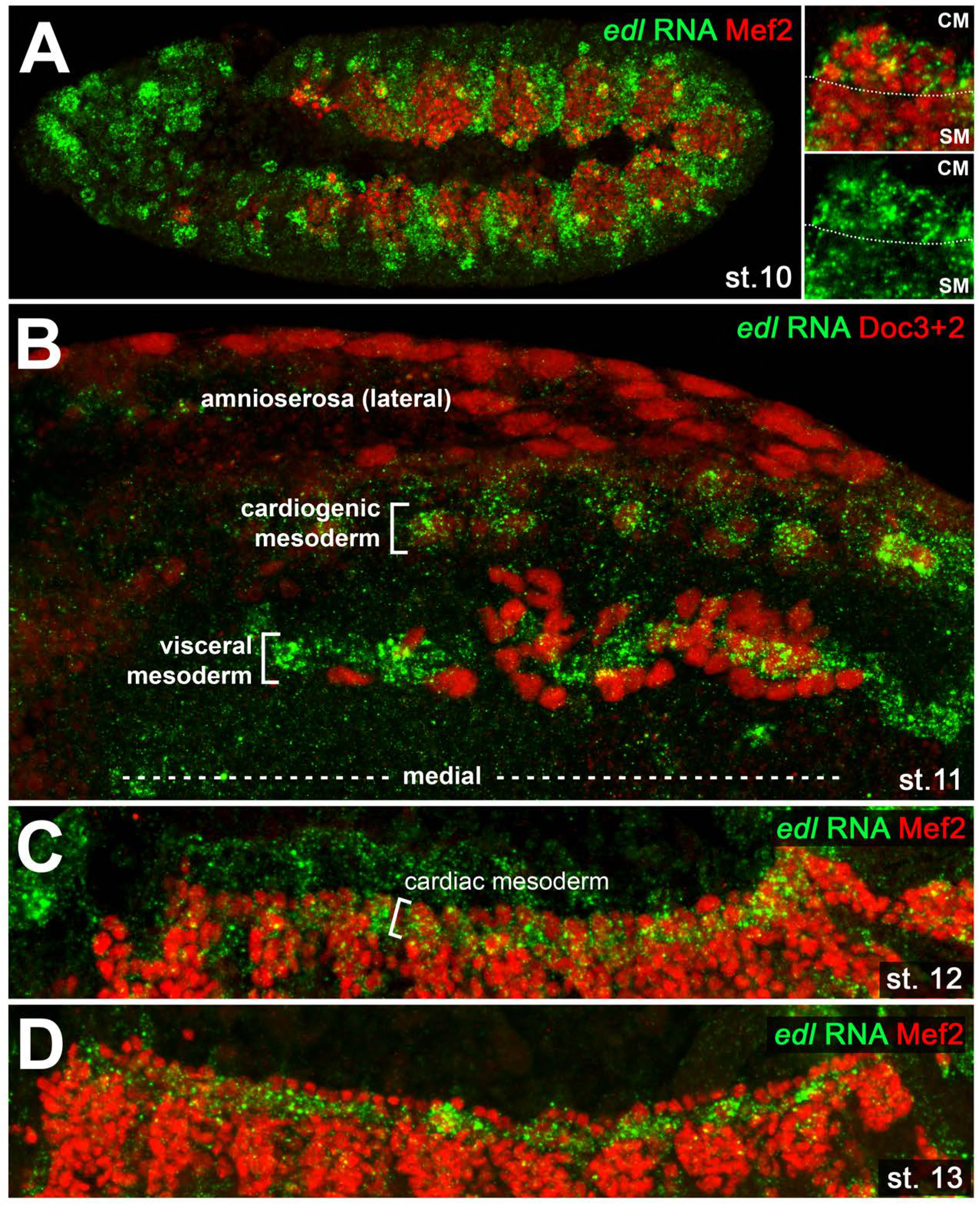
Cardiac *edl* expression. Detection of *edl* RNA in wild type embryos co-stained against Mef2 (A,C,D; lateral views) or Doc (B; dorsal view). (A) Stage 10 embryo showing strong *edl* expression in numerous ectodermal and mesodermal tissues including the Mef2-positive areas of the early cardiogenic mesoderm (CM) and parts of the somatic mesoderm (SM). (B) Stage 11 embryo in which *edl* RNA is strongly expressed in Doc^+^ cardiogenic clusters. High expression is also seen in the band of trunk visceral mesoderm founders, but not in adjacent migrating longitudinal visceral muscle founders (also Doc^+^). (C) Cardiac *edl* expression persists during germ band retraction. (D) Thereafter, it fades in the cardioblasts but continues to be expressed in the pericardial region.

**Figure 4 - source data 1. Quantification of cardioblasts in various genotypes affecting Edl, Pnt or Yan/Aop activities.** data file: fig4-data1.xlsx

**Figure 5 - Figure supplement 1.**
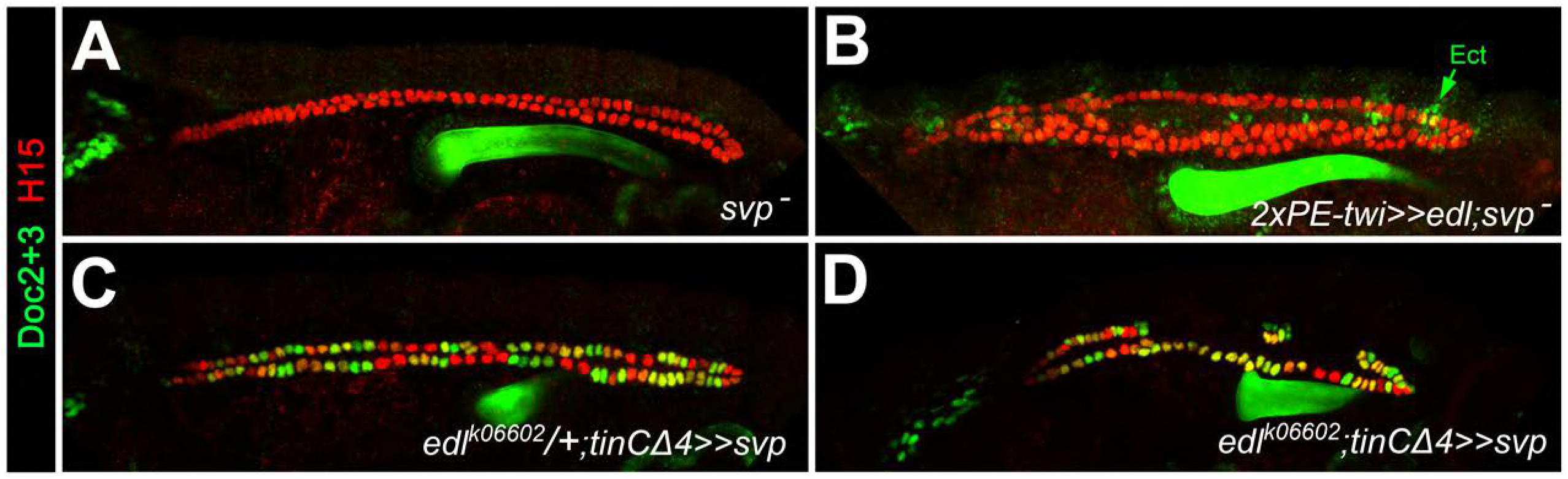
Epistatic relationship between *edl* and *svp*. Embryos stained for H15 and Doc as the wild-type control in Figure 1A. (A) Loss of *Doc* expression in homozygous *svp*^*AE127*^ mutants. (B) All CBs remain Doc-negative upon pan-mesodermal *edl* overexpression in the *svp*^*AE127*^ mutant background. (C) Overexpression of *svp* in CBs leads to ectopic *Doc* expression. Sporadic reduction in H15 expression (green CBs) may result from repression of *tin*, since similar H15 reductions were observed in CB-specific *tin*-mutants (Figure 3H). (D) In the absence of *edl*, forced expression of *svp* with the same driver also expands the population of Doc^+^ CBs. This suggests that *svp* is epistatic to *edl* during of CB patterning.

**Figure 6 - Figure supplement 1.**
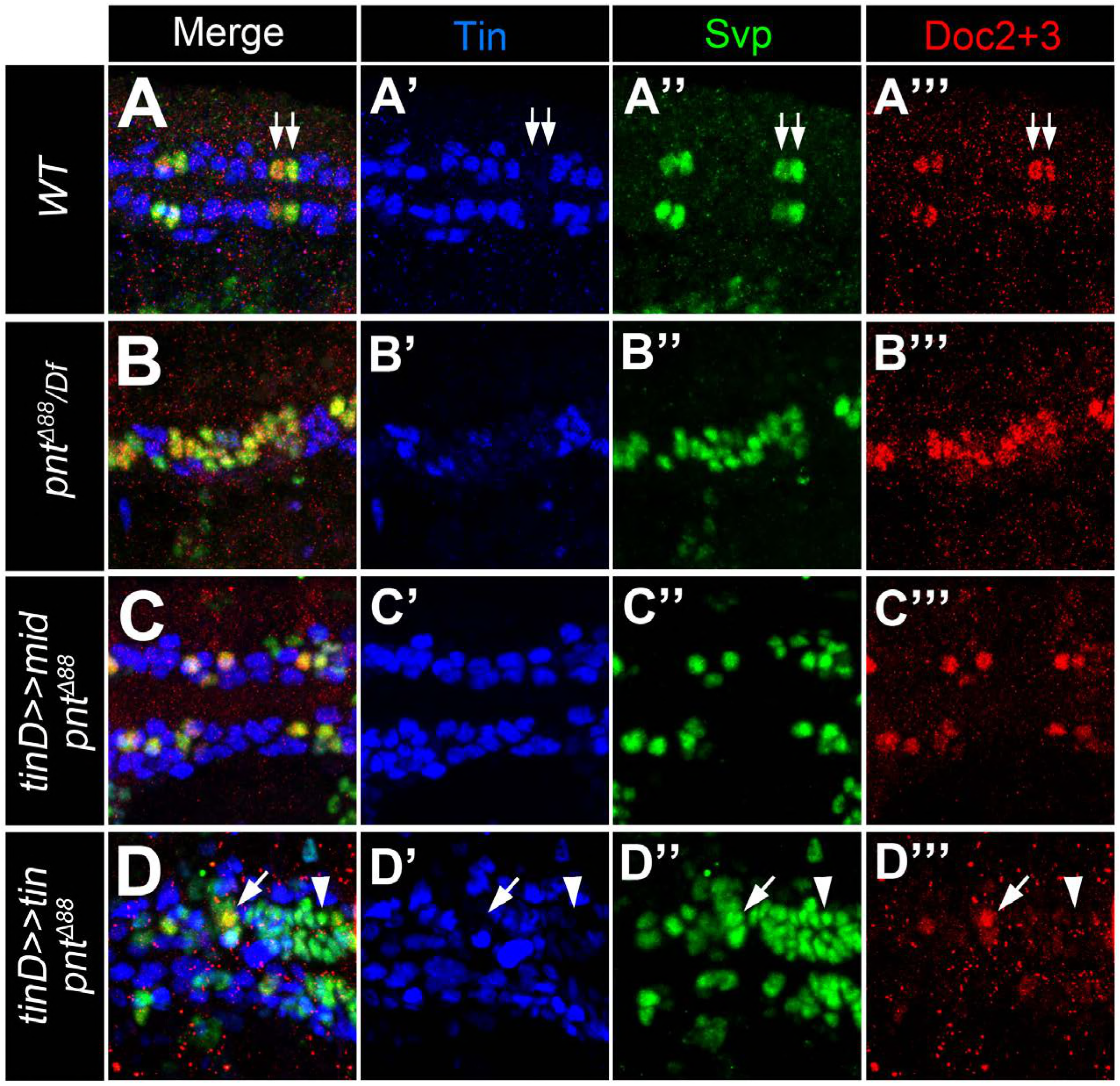
Additional data supporting *mid* function in gCBs. Stainings with antibodies against Svp (green), Doc2+3 (red) and Tin (blue). Shown are representative regions of the developing heart in stage 15-16 embryos (merged and single channels as indicated). (A) Control embryo showing co-expression of Svp and Doc in the Tin-negative oCBs (arrows). (B) In amorphic *pnt* mutants (this example: *pnt*^*Δ88*^/*Df(3R)Exel9012*) expression of Svp expands simultaneously with Doc. (C) Overexpressing *mid* in a *pnt* null background in the dorsal mesoderm via *tinD*-GAL4 largely reverts the expansion of both Doc and Svp. (D) By contrast, overexpressing *tin* with the same driver in *pnt* mutants has a repressive effect on Doc but not Svp expression (arrowheads).

**Figure 7 - Figure supplement 1.**
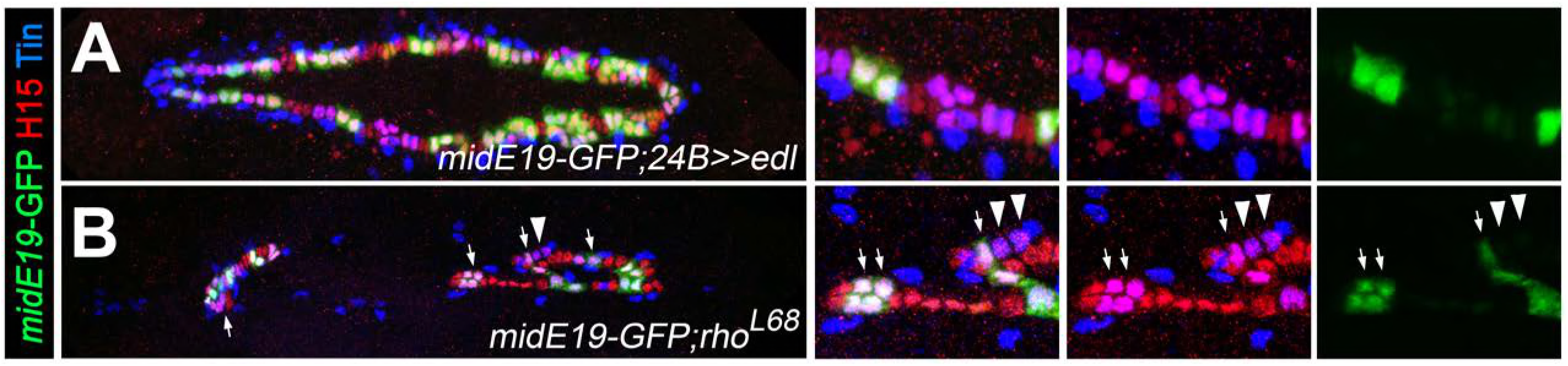
Further analysis of *midline* regulation. Stainings for GFP (green), H15 (red) and Tin (blue) in stage 16 embryos carrying the *midE19-GFP* reporter as the control in Figure 7A. (A) Mesodermal overexpression of *edl* via *how^24B^-*GAL4 leads to a loss of *midE19*-GFP in many gCBs (purple). (B) Loss of *rho* function leads to a complete loss of GFP in some of the retained gCBs (arrowheads) and a level reduction in others (arrows). In comparison to *pnt* mutants (Figure 7B), a higher fraction of gCBs retains substantial GFP expression indicating additional, *rho-*independent inputs upstream of Pnt.

